# Genome-wide screen identifies novel factors for surface protein cross-wall trafficking and cell envelope homeostasis in *Staphylococcus aureus*

**DOI:** 10.1101/2025.11.03.686427

**Authors:** Salvatore J. Scaffidi, Ran Zhang, Yaosheng Jia, Wenqi Yu

## Abstract

Cell wall anchored surface proteins are integral components of the Gram-positive bacterial cell envelope and are vital for bacterial survival in different environmental niches. The trafficking of many surface proteins carrying a YSIRK/G-S signal peptide is synchronized with cell envelope biogenesis during cell division, whereby YSIRK proteins traffic to the septal membrane and anchor to septal peptidoglycan (cross-wall). Previous work demonstrated that LtaS-mediated lipoteichoic acid (LTA) synthesis restricts YSIRK proteins septal trafficking. Here we did a comprehensive immunofluorescence microscopy screen of the entire *S. aureus* Nebraska Transposon Mutant Library (NTML) for additional factors regulating cross-wall trafficking of staphylococcal protein A (SpA), an archetype of YSIRK proteins. We characterized the top nine major hits that drastically diminished SpA cross-wall localization, including *ypfP* and *ltaA* (LTA glycolipid anchor synthesis genes), *lcpB (*LytR-CpsA-Psr family protein), *mprF* (lysyl-phosphatidylglycerol synthase), *lytH* (cell wall hydrolase), *scdA* (nitrite reductase)*, yjbH* (protease adaptor protein )*, cbiO* (cobalt transporter) and *SAUSA300_2311* (LytTR regulatory system) along with Δ*tagO* (wall teichoic acid synthesis). Interestingly, unlike the *ltaS* mutant that delocalizes SpA at both the septal membrane and peptidoglycan (PG) layer, all the hits only delocalized SpA at the PG layer, suggesting that these mutants affect the late-stage SpA trafficking. In addition, mutants of *lcpB, yjbH, cbiO* and *2311* exhibit both transcriptional and spatial regulation. All the hits showed defects in cell cycle, cell morphology and spatially dysregulated PG synthesis. The shared phenotypes among the mutants suggest that impaired PG homeostasis and cell cycle defects are the mechanisms underlying dysregulated SpA localization. Overall, this work not only expands our understanding of YSIRK protein cross-wall trafficking but also identifies new leads that have a broader impact on the dynamics of cell cycle and cell envelope homeostasis.

**Importance:** Surface proteins of gram-positive bacteria are key virulence factors in human pathogen *S. aureus.* Most surface proteins carry a YSIRK/G-S type signal peptide that promotes cross-wall trafficking and attachment to septal cell wall during cell division. This study identified several new factors regulating this process through a comprehensive screen. The mutants identified here display dysregulated cell wall synthesis along with cell cycle defects. The results provide new insight into virulence factor trafficking and cell envelope homeostasis, which lays the foundation for development of new antimicrobial therapeutics.

## Introduction

Surface proteins of gram-positive bacteria are covalently attached to cell wall peptidoglycan (PG) and play key roles in bacterial survival and adaptation to harsh environments. For pathogenic bacteria, these proteins are essential in adhesion to host tissues, colonization, and immune evasion, making them as promising vaccine candidates (1–3). A highly conserved biochemical sorting pathway mediates the trafficking of surface proteins from cytosol to bacterial cell surface (4). The sorting pathway is comprised of three main stages: secretion, anchoring and incorporation. Surface protein precursors contain an N-terminal signal peptide and a C-terminal cell wall sorting motif. The N-terminal signal peptide is recognized by SecA and translocated through the SecYEG translocon across the cytoplasmic membrane (5–7). The signal peptide is cleaved off by the type I signal peptidase (SpsB in *S. aureus*) upon membrane translocation (8–10). The membrane-bound transpeptidase sortase A (SrtA) recognizes and cleaves the C-terminal LPXTG cell wall sorting motif and covalently anchors the pre-protein to PG precursor lipid II (11). Lastly, the lipid II-protein complex is incorporated into mature PG via transglycosylation and transpeptidation during cell wall biosynthesis (12–14).

Remarkably, many gram-positive surface proteins possess a conserved YSIRK/G-S motif within the signal peptide (7, 15, 16). In their pioneering paper from 1962, Cole and Hahn demonstrated that newly anchored M protein appeared at the cross-wall in *Streptococcus pyogenes,* and M protein was used to label the site of new cell wall synthesis (17). The cross-wall localization of M protein was attributed to its YSIRK/G-S signal peptide (18). The same phenotype was found in *S. aureus* (5). It is proposed that surface protein precursors with YSIRK signal peptide are secreted at the septum and anchored to cross-wall during cell division (5, 6). The cross-wall anchored proteins eventually distribute all over the cell surface after consecutive rounds of cell division and separation (19). Thus, the dynamic YSIRK protein trafficking is coupled with new PG synthesis, cell division and cell cycle.

The gram-positive bacterial cell envelope is composed of a single cytoplasmic membrane and a thick cell wall layer (12, 20, 21). The cell wall is composed of cross-linked glycan chains, which acts as a scaffold for proteins and teichoic acids (20–26). Wall teichoic acids (WTA) are anchored directly to PG and are composed primarily of polyribitol phosphate chains, which extrude through the cell surface (23, 27–29). Lipoteichoic acids (LTA) are glycerol-phosphate polymers tethered to the cytoplasmic membrane via a glycolipid anchor and extend through the membrane-wall interface underneath the PG matrix (24, 30). In the cytoplasm, YpfP synthesizes diglucosyl-diacylglycerol (Glc2-DAG) - the glycolipid anchor, which is translocated across the membrane by LtaA (31, 32). LtaS synthesizes the LTA polymer by repeatedly transferring glycerol phosphate moieties from phosphatidylglycerol to Glc2-DAG (32, 33). Previous work demonstrates that YSIRK protein trafficking is synchronized with LTA synthesis: LtaS-mediated LTA synthesis spatially restricts staphylococcal protein A (SpA), an archetype of YSIRK proteins in *S. aureus* (6, 30). Mutants of *ltaS* diminishes SpA localization at the septal membrane and cross-wall (6, 30). Mature LTA localizes at the peripheral cell membrane, providing a restriction mechanism for SpA septal localization (30). The signal peptidase SpsB coordinately processes SpA pre-proteins and LtaS at the septal membrane, which is essential for SpA septal secretion (10). D-alanylation of teichoic acids as well as mutants of *gdpP* modulate SpA cross-wall localization but did not affect SpA septal secretion (30, 34). Recent work from a screen of temperature sensitive mutants identified *secA(ts*) allele as well as mutations in the *secG* and *pepV* genes (35). PepV interacts with SecA and SpA pre-proteins and exhibits a mutually inhibitory effect on the stability of SpA pre-proteins (35). A screen from an arranged transposon library identified mutants of *sscB* and *sscE* (encoding staphylococcal surface carbohydrate synthesis gene B and E) impaired SpA cross-wall localization (36), for which the mechanism is unclear. Furthermore, staphylococcal major autolysin *atlA* was implicated in SpA surface display and release (37).

The above research indicates that the septal trafficking of YSIRK proteins is closely connected to the biogenesis of cell wall and other cell envelope structures. Multiple factors may modulate YSIRK protein trafficking at different stages. Inspired by the work of Cole and Hahn in 1962, where M protein was used to label new cell wall synthesis sites, we did a comprehensive screen of the Nebraska Transposon Mutant Library (NTML) with the aim of identifying additional factors that spatially regulate YSIRK protein trafficking and cell envelope biogenesis. We selected nine major hits that drastically delocalized SpA and performed detailed characterization. These mutants displayed strong morphological defects, spatially altered PG synthesis and autolysis. Upon completion of this screen, we have defined previously uncharacterized genes, along with new functions for those known genes involved in cell envelope biogenesis. This work not only expands our understanding of YSIRK surface protein septal trafficking but also identifies new leads that broadly impact cell envelope homeostasis and metabolism during cell division and cell growth.

## Results

### Immunofluorescence microscopy screen identified mutants deficient in localizing SpA at cross-wall

We previously established SpA immunofluorescence microscopy (IF) protocols to examine SpA localization at the septal membrane (membrane IF) and cross-wall (cross-wall IF) of *S. aureus* (38, 39). Here, we standardized the cross-wall IF protocol to screen all mutants of the NTML (**Fig. 1A**). Briefly, bacterial cells were grown to mid-exponential phase and trypsin-treated to remove pre-existing SpA from the cell surface. Trypsin-treated cells were further grown for 20 minutes (equal to one round of cell division) in fresh media containing trypsin inhibitor, which allows deposition of newly synthesized SpA to the cell wall. The cells were fixed immediately, stained with SpA first and secondary antibodies and visualized by fluorescence microscopy. Cells were also counter-stained with DNA and membrane dye. As a control, we replaced the SpA signal peptide (SP*_spa_*) with a non-YSIRK signal peptide (SP*_sasD_*) in fusion with mature SpA (SP_SasD_-SpA). The SP_SasD_-SpA translational fusion is expressed under *spa* native promoter at an ectopic locus in the chromosome of JE2Δ*spa::Tn* (**Fig. 1B**).

**Fig. 1.**
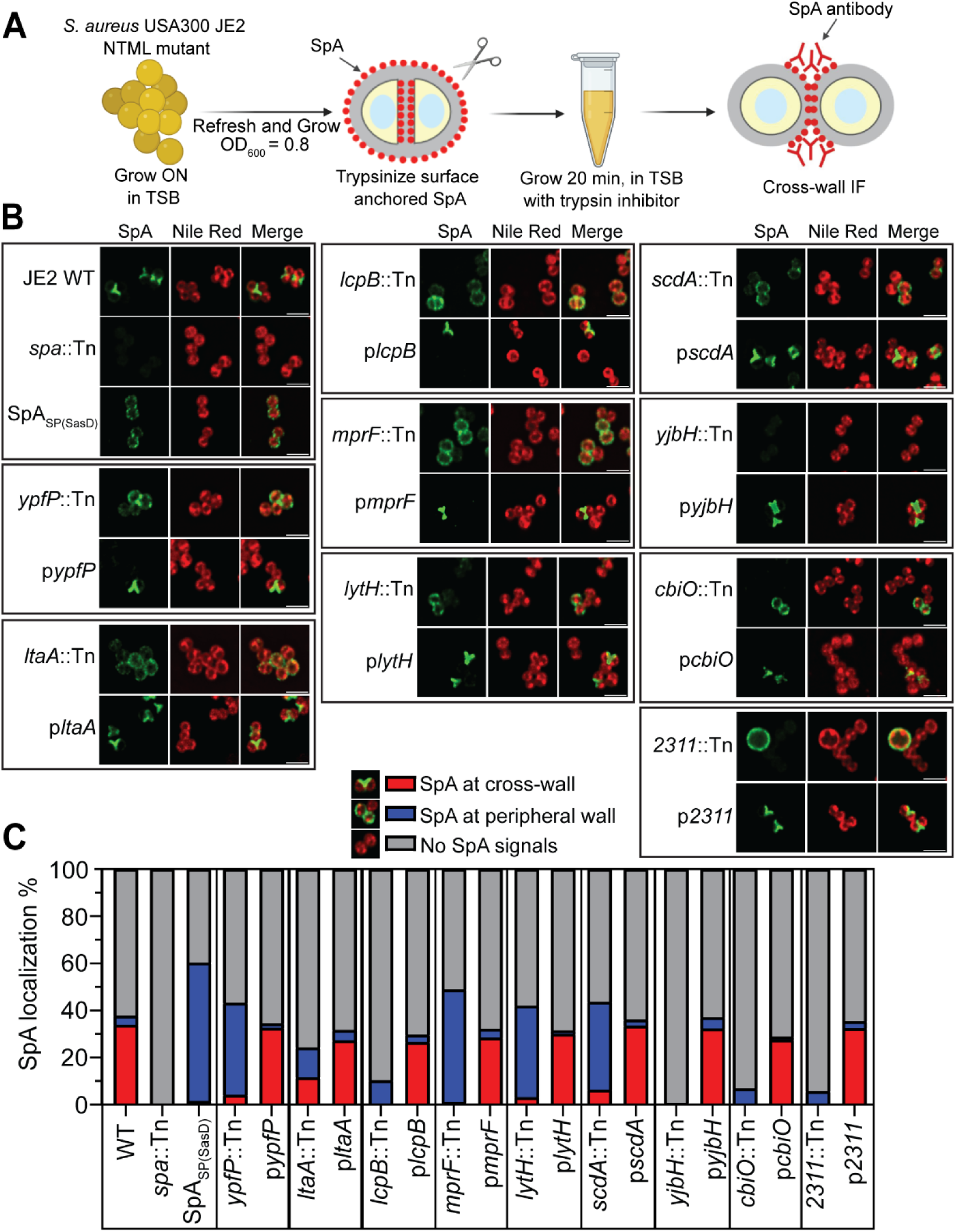
Genome-wide screen identifies mutants that diminish SpA cross-wall localization. A) Outline of cross-wall immunofluorescence (IF) microscopy screen of the Nebraska transposon mutant library (NTML). B) Localization of de novo SpA on staphylococcal cell surface revealed by cross-wall IF microscopy in NTML screen major hits. Green fluorescence shows SpA and Nile red stains cell membrane (red). Scale bar = 2 µm. C) Quantification of SpA localization at the cross-wall (red bar), peripheral wall (blue bar) or no detectable SpA signals (gray bar) from the images represented in panel B. Mean values and standard deviation are shown in Table S3.

The entire NTML contains 1,952 mutants in USA300 JE2, a community acquired methicillin-resistant *S. aureus* (CA-MRSA) strain (40). Each mutant possesses a single insertion of the *bursa aurealis* mariner-based transposon (Tn) in the coding sequence of one nonessential gene (40). We screened a total of 1,914 mutants because some strains were missing in our collection. Each mutant was examined alongside controls of JE2 wild type (WT), a *spa* transposon mutant (JE2Δ*spa*::Tn) and JE2 *spa*::Tn expressing SP_SasD_-SpA (**Fig. 1B**). While JE2 WT exhibits the typical SpA localization at the cross-wall between two daughter cells (visualized as ‘Y’ or ‘X’ shape or a line), SP_SasD_-SpA displays weaker SpA signals and more homogenous and circumferential localization all over the cell wall (**Fig. 1B**). JE2Δ*spa*::Tn is a negative control with no SpA signals. The phenotype of each NTML mutant was compared with the controls and scored based on the overall SpA fluorescence signal intensity and SpA localization. The mis-localization of SpA is defined by two criteria: (a) the frequency of cross-wall localized SpA, which is quantified by the numbers of cells showing cross-wall SpA in a cell population (**Fig. 1C**); (b) the signal intensity of cross-wall localized SpA, which is quantified by the ratio of cross-wall SpA fluorescence intensity versus peripheral wall SpA fluorescence intensity **(Fig. S1).** Our screen identified mutants that are known to regulate SpA production and localization, such as *srtA* (11)*, sagB* (41)*, xdrA* (42)*, codY* (43), and *sarA* (44), which validates the robust nature of our screen.

It is known that the expression of *spa* is under the control of multiple transcriptional regulators at its extended promoter sequence (45). Either over-expression or under-expression of *spa* can affect SpA localization, which complicates the interpretation of our screening results. To distinguish transcriptional versus spatial regulation, all the major hits were transduced to a different strain WYL478 (RN4220Δ*spa* pCL55-P*_itet_*-*spa*), where the native promoter of *spa* is replaced by an ATc-inducible promoter (**Fig. 2**). This experiment eliminated mutants that only affect *spa* expression, but not SpA localization, which we did not follow up. We continued to characterize the top nine major hits that showed severe SpA mis-localization as discussed below.

**Fig. 2.**
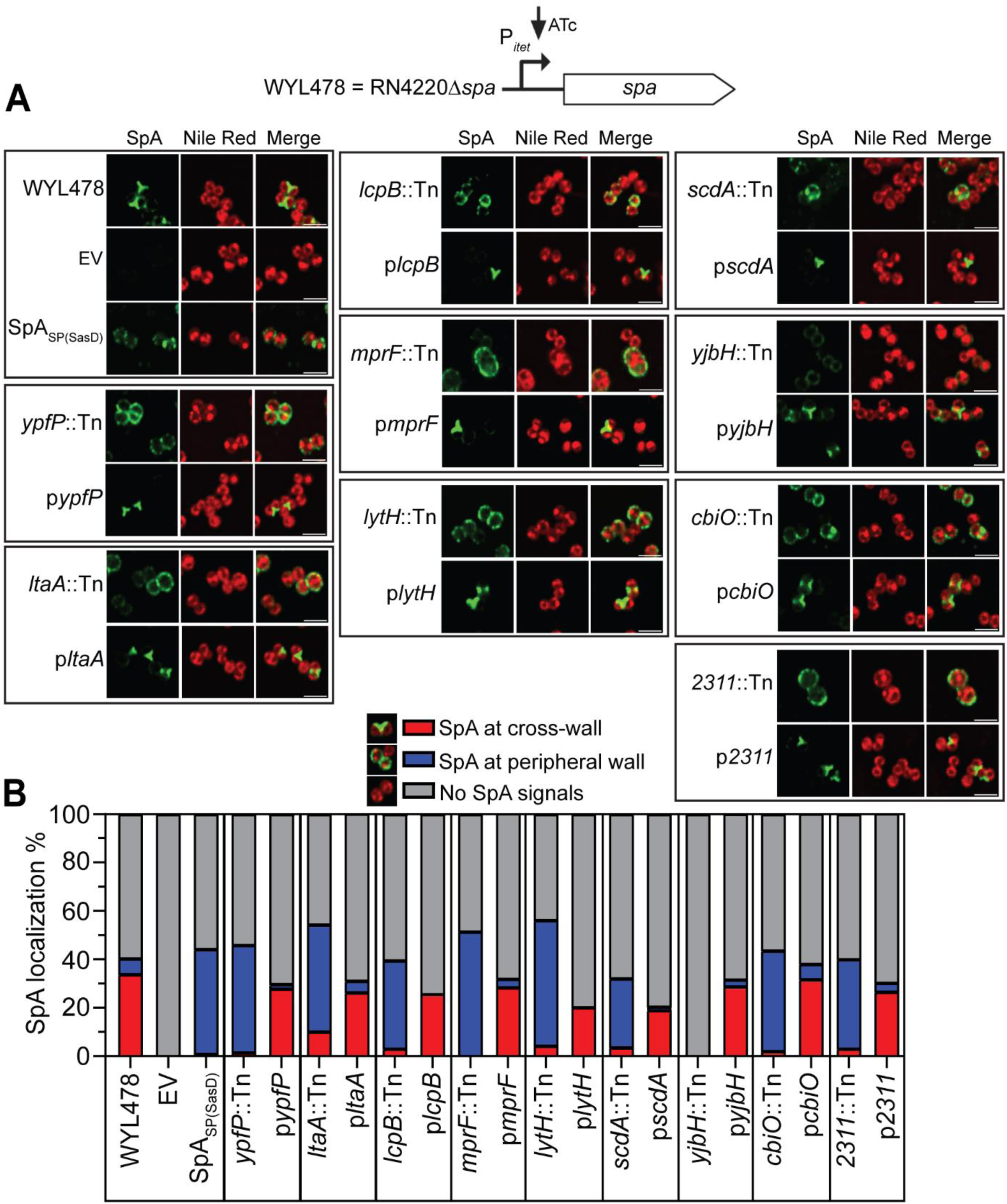
Cross-wall IF of mutants transduced to WYL478. A) Cross-wall IF images showing SpA localization in transductant mutants in WYL478 background, grouped with respective complement strain. B) Quantification of SpA localization at the cross-wall (red bar), peripheral wall (blue bar) or no detectable SpA signals (gray bar) from the images represented in panel B. Mean values and standard deviation are shown in Table S3.

### Mutants of *ypfP, ltaA, mprF, lytH, scdA* diminish SpA cross-wall localization without affecting *spa* expression, whereas mutants of *lcpB, yjbH, cbiO, 2311* affect both *spa* expression and SpA localization

The top nine major hits are mutants of *ypfP, ltaA, mprF, lytH, lcpB, scdA, yjbH, cbiO, 2311*. YpfP and LtaA are involved in LTA glycolipid anchor synthesis as mentioned above (46). MprF stands for staphylococcal multiple peptide resistance factor, which synthesizes lysyl-phosphatidylglycerol by adding a positively charged lysyl group onto phosphatidylglycerol (47, 48). LytH is a membrane-bound cell wall hydrolase that removes stem peptides from uncross-linked PG (49). LcpB belongs to the LytR-CpsA-Psr (LCP) family proteins implicated in attaching WTA to PG (50). ScdA was initially found to be a determinant of proper cell division and morphogenesis and has recently been shown to have nitrite reductase activity converting nitrite to nitric oxide (51, 52). YjbH is a ClpXP protease adaptor protein that binds to the stress-inducing regulator Spx and enhances its degradation (53). The last two hits *cbiO* (*SAUSA300_2176*) and *SAUSA300_2311* are uncharacterized genes. CbiO is denoted as cobalt transporter ATP-binding subunit or energy-coupling factor transporter ATPase (EcfA1). *SAUSA300_2311* is in an operon with its upstream gene *SAUSA300_2310,* which is homologous to a LytTR regulatory systems in *Streptococcus mutans* (54), whose function is not known in *S. aureus*.

Figure 1 summarizes the cross-wall IF screen results of the major hits. Compared to the WT where SpA signals intensified at the cross-wall, all the mutants drastically diminished SpA cross-wall signals. SpA appeared to be circumferentially distributed all over the cell wall in these mutants similar to the SP_SasD_-SpA control (Fig. 1B). In addition, *yjbH::Tn* showed no detectable SpA signals. Mutant of *2311* showed heterogeneous cell sizes; SpA showed circumferential distribution in only the enlarged cells while the rest of cells did not display SpA signals. Each mutant was complemented by expressing the gene in a multi-copy plasmid. Quantification of SpA localization showed that mutants of *ypfP, ltaA, mprF, lytH* and *scdA* significantly decreased SpA cross-wall localization and increased SpA peripheral wall localization (Fig. 1C). Quantification of SpA signal fluorescence intensity (ratio of cross-wall versus peripheral wall) confirmed circumferential mis-localization of SpA particularly in these five mutants **(Fig. S1)**. Mutants of *lcpB, yjbH, cbiO* and *2311* had an increased cell population without SpA signal (Fig. 1C).

When the mutants were examined in WYL478 strain background where *spa* is expressed by an ATc-inducible promoter, SpA overall signals in the mutants of *lcpB, cbiO* and *2311* were restored to WT level while SpA remained mis-localized (Fig. 2). SpA signals were increased but still weak in *yjbH::Tn* (Fig. 2). All the other mis-localization mutants (*ypfP, ltaA, mprF, lytH* and *scdA*) showed similar SpA mis-localization in WYL478 as in JE2, indicating that the mis-localization phenotype is reproducible independent of strain backgrounds.

To further examine if the mutants affected SpA overall protein abundance, the proteins from cell pellet (P) and supernatant (S) of all mutants from both JE2 and WYL478 background were collected and examined by SpA immunoblots (Fig. 3). Consistent with microscopy observation, SpA protein level was significantly reduced in the mutants of *lcpB, yjbH, cbiO* and *2311* in JE2, and was restored in WYL478.

**Fig. 3.**
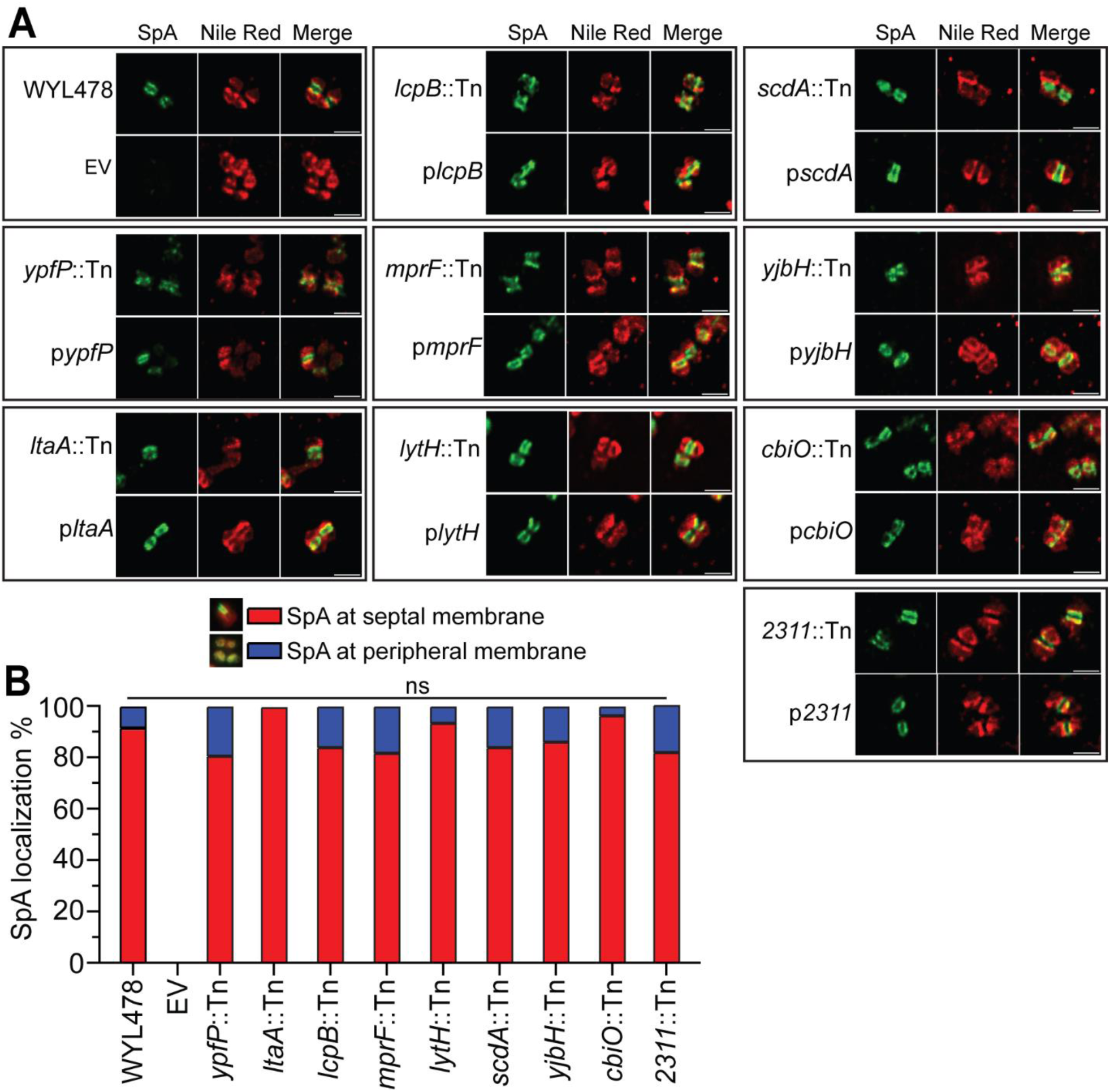
Membrane IF of WYL478 tranductant mutants. A) Membrane IF microscopy images showing SpA localization at the septal membrane in WYL478 transductants, grouped with respective complement strain. B) Quantification of SpA localization at the septal membrane (red bar) and peripheral membrane (blue bar) from the images represented in panel A. ns = no significant difference.

Taken together, these results indicate that SpA is transcriptionally and spatially regulated in the mutants of *lcpB, yjbH, cbiO* and *2311,* whereas mutants of *ypfP, ltaA, mprF, lytH* and *scdA* dysregulate SpA localization.

### None of the mutants affected SpA septal membrane localization

To further investigate the effect of these mutants on SpA localization, we examined SpA localization at the septal membrane by a membrane-IF method that we previously established (38). Briefly, trypsinized cells were fixed and the cell wall was digested with the cell wall hydrolase lysostaphin; the resulting protoplasts were fixed and stained with SpA primary and secondary antibodies. We examined all the WYL478 mutants along with complementation strains. Interestingly, SpA was dominantly localized at the septal membrane in all the mutants similar to the WT (Fig. 4A). Quantification of cells displaying SpA at the septal membrane versus the peripheral membrane yielded no significant difference across all strains (Fig. 4B). The septal membrane localization of SpA revealed by membrane-IF is dependent on the membrane transpeptidase SrtA (30). The results that SpA septal membrane localization was not affected suggest that SrtA-mediated anchoring process is not affected in these mutants. Instead, all the mutants likely affect the late-stage SpA trafficking post SrtA-mediated anchoring.

**Fig. 4.**
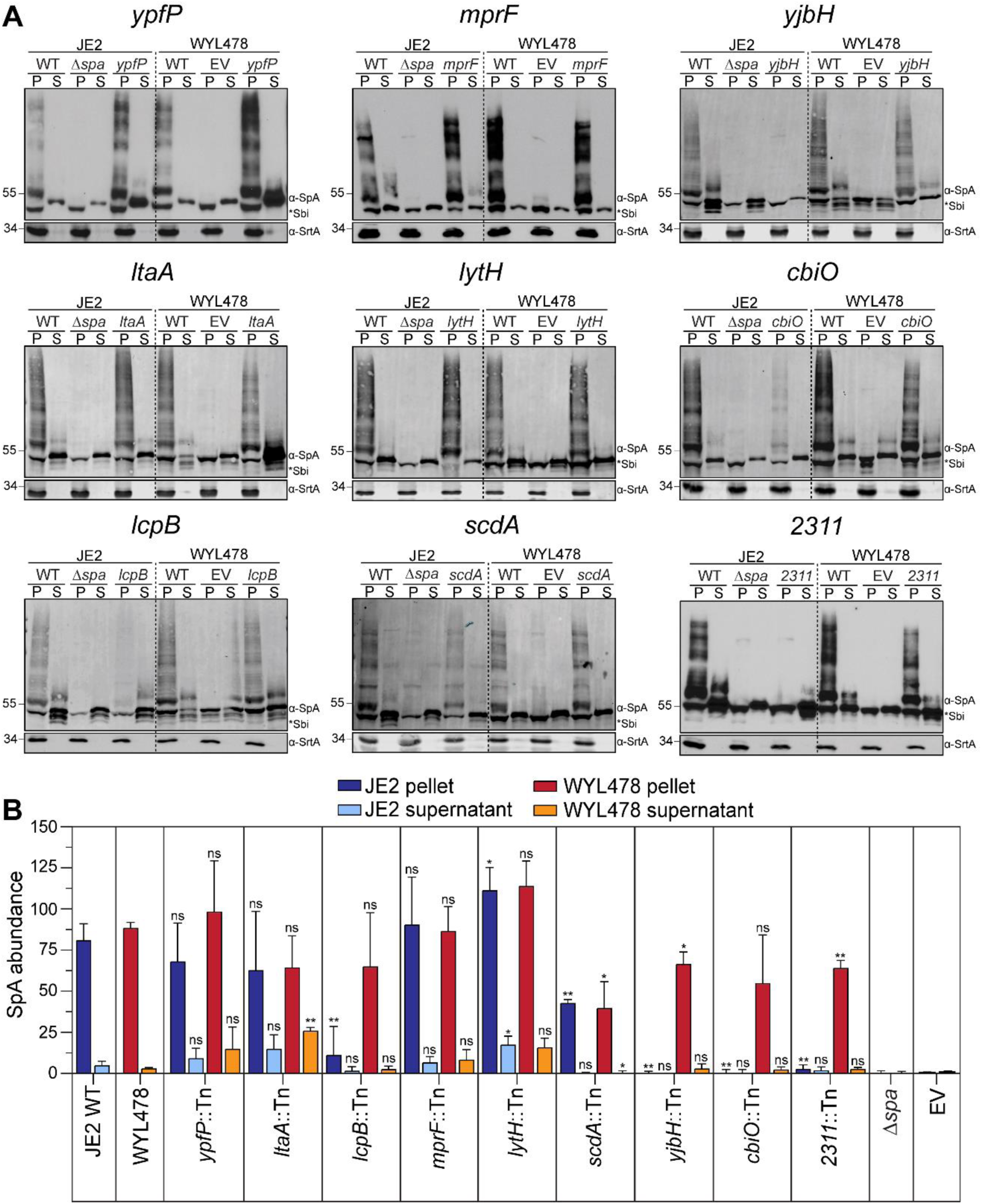
SpA immunoblots of the mutants in JE2 and WYL478. A) Western blot analysis of SpA in cell pellet (P) and supernatant (S). Sortase A (SrtA) blot is a loading control. All the blots were repeated three times and one representative blot is shown. B) Quantification of SpA band intensity divided by SrtA loading control band to determine normalized SpA abundance. Unpaired *t*-test with Welch’s correction was performed for statistical analysis: \**P* < 0.05, \*\**P* < 0.005, \*\*\**P* < 0.0005, \*\*\*\**P* < 0.0001.

### Characterizing cell cycle and morphological defects in the mutants

Once SpA is anchored to PG by SrtA, its localization is coupled with the dynamics of cell wall homeostasis including cell wall synthesis and turnover throughout the cell cycle. Thus, in the next series of experiments, we examined the cell cycle, cell wall synthesis and autolysis profiles of the major hits.

To examine cell cycle and cell morphology, mutant cells were stained with fluorescently labeled vancomycin (Van-FL) that binds to uncrosslinked D-Ala-D-Ala residues in the PG. Mutants of *ypfP, ltaA, mprF* and *2311* exhibited severe morphological defects shown as abnormal septum formation, heterogeneous cell size and irregular cell clusters formation (Fig. 5A**, arrowheads)**. Quantification of cell size indicated that mutants of *ypfP, ltaA, lcpB, mprF, scdA* and *2311* had significantly enlarged cell size, whereas mutants of *yjbH* and *cbiO* showed decreased cell size (Fig. 5C). *2311::Tn* exhibited striking heterogeneity in cell size, with some very enlarged cells showing doubled cell size of WT cells. To analyze cell cycle progression, we quantified the three cell cycle phases as defined previously (10, 55). Cells in phase 1 display no septum, whereby they have completed their previous division and have not yet begun the next round. Cells in phase 2 represent those with partial septum formation, which have initiated cell division but not yet completed septum constriction. Lastly, cells in phase 3 have a complete septum at the mid-cell. Compared to WT, all the mutants exhibited altered cell cycle dynamics with notably increased phase 2 cells (Fig. 5B). Mutants of *ypfP*, *ltaA*, *mprF*, *scdA*, *yjbH* and *2311* displayed roughly a two-fold increase in phase 2 cells. The defects in cell cycle and cell morphology were restored to WT level with complementation. We did the same experiment with the transductant mutants in WYL478, and the same results were obtained **(Fig. S2)**. Collectively, these results indicate that the mutants are impaired in cell cycle progression with deficiency in cell septation and division.

**Fig. 5.**
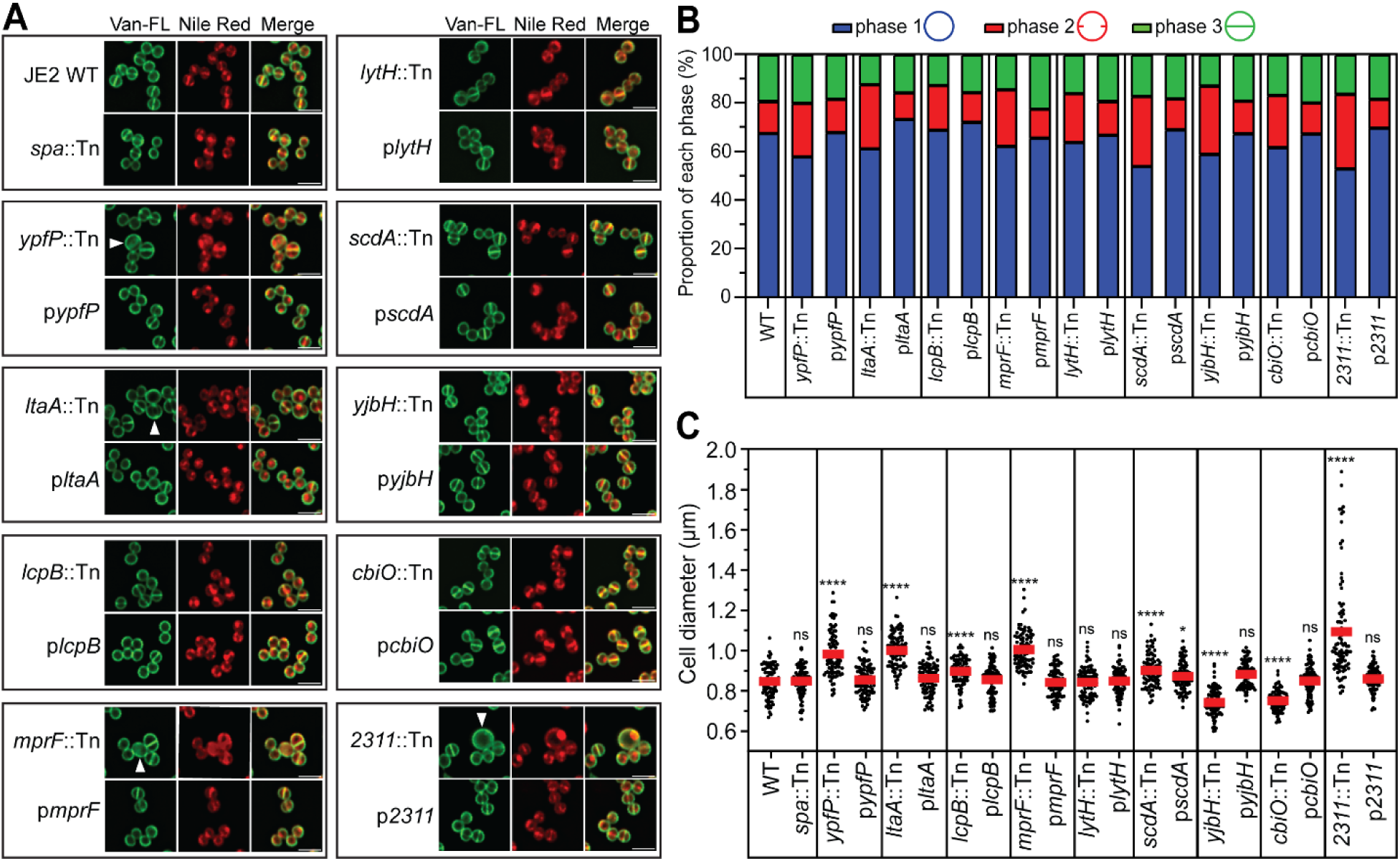
Fluorescent vancomycin (Van-FL) labeling reveals defects in cell morphology and cell cycle in the NTML mutants. A) Fluorescence images showing the cell wall (Van-FL) and membrane (Nile red). White arrowheads indicate cells with aberrant morphology compared to JE2 WT. B) Cell cycle quantification reveals septation defects in screening mutants. Van-FL images were quantified to determine the proportion of cells in phase 1 (no septum, blue), phase 2 (partial septation, red) or phase 3 (completed septum, green). C) Quantification of cell diameters. Quantification of cell diameter across mid-cell using the Van-FL images represented in panel A, n = 90 cells. Each mutant and complementation strain is compared to the JE2 WT. Unpaired *t*-test with Welch’s correction was performed for statistical analysis: \**P* < 0.05; \*\**P* < 0.005; \*\*\**P* < 0.0005; \*\*\*\**P* < 0.0001.

### The dynamic PG synthesis is spatially dysregulated in the mutants

It is known that new PG synthesis occurs at the septum (56), and the next round of cell division occurs perpendicular to the previous one in staphylococci (57). As a result of successive rounds of cell division, the previously synthesized PG accumulates at the cell periphery. This dynamic PG synthesis and its spatial distribution can be monitored by sequential labeling of fluorescent D-amino acids (FDAAs) (10). All mutant cells were sequentially incubated with three different FDAAs (blue HADA, red RADA, and green OGDA) to track PG incorporation along the cell cycle. As shown in Fig. 6, the green OGDA labels the new septum, the red RADA labels the cross-wall (old septum) and the blue HADA labels the old peripheral wall in WT cells. In comparison to JE2 WT, all mutants displayed various FDAA mis-incorporation patterns. Those mutants with strong morphological defects (*ypfP, ltaA, lcpB, mprF and 2311*) showed severe defects in FDAA incorporation, where the dynamic distribution of FDAAs was drastically altered (Fig. 6A**, arrowheads)**. Mutants of *lytH, scdA, yjbH* and *cbiO* showed delayed FDAA incorporation, where two or even three FDAAs colocalized (Fig. 6A**, arrowheads).** The percentage of abnormal FDAA incorporations was quantified (Fig. 6B). FDAA staining of the WYL478 transductants revealed similar results **(Fig. S3)**. Overall, all mutants dysregulate the spatial dynamics of PG synthesis.

**Fig. 6.**
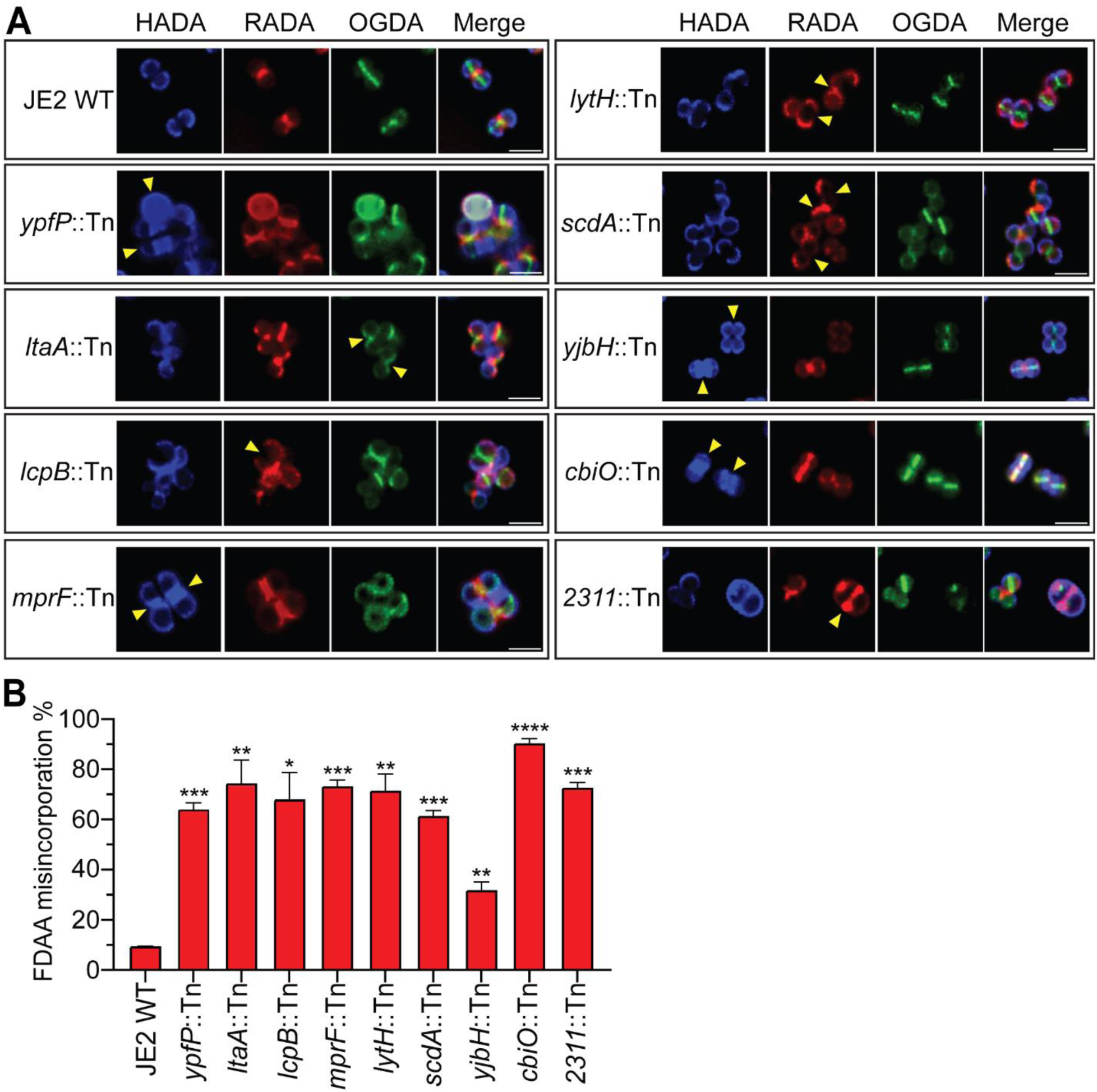
Sequential Fluorescent D-amino acids (FDAAs) incorporation reveals spatially dysregulated cell wall biosynthesis in NTML mutants. A) Fluorescence images of cells labeled with FDAAs: HADA (blue, first), RADA (red, second) and OGDA (green, third), yellow arrows indicate aberrant FDAA localization. B) Quantification of FDAA misincorporation. Unpaired t-test with Welch’s correction was performed for statistical analysis: \**P* < 0.05, \*\**P* < 0.005, \*\*\**P* < 0.0005, \*\*\*\**P* < 0.0001. Representative images and quantification are from three independent experiments.

### Altered autolysis profiles in the mutants

Cell wall homeostasis is modulated by PG hydrolases (autolysins). To assess the activity of autolysins and the susceptibility of the mutants to autolysis, we performed the Triton X-100 mediated autolysis assay. Triton X-100 is a non-ionic detergent which stimulates autolysin activity under static growth conditions in *S. aureus* (58). In WT cells, Triton X-100 induced gradual lysis over time, whereas a mutant lacking the major staphylococcal autolysin AtlA (*altA*::Tn) showed no lysis (Fig. 7). Compared to the controls of JE2 WT and *atlA*::Tn, the mutants of *ypfP, ltaA, lcpB*, *mprF* and *2311* showed significantly increased autolysis, the mutant of *cbiO* showed decreased autolysis and the mutants of *lytH, scdA* and *yjbH* did not show significant change (Fig.7). The increased susceptibility to autolysis seemed to correlate with severe morphological defects and FDAA misincorporation in the mutants of *ypfP, ltaA, lcpB*, *mprF* and *2311.* While LytH is a cell wall hydrolase, *lytH::Tn* did not show apparent change in autolysis. The altered autolysis in the mutants of *ypfP, ltaA, lcpB*, *mprF*, *2311 and cbiO* are likely to due to their impaired cell wall structure, which in turn modulates SpA localization.

**Fig. 7.**
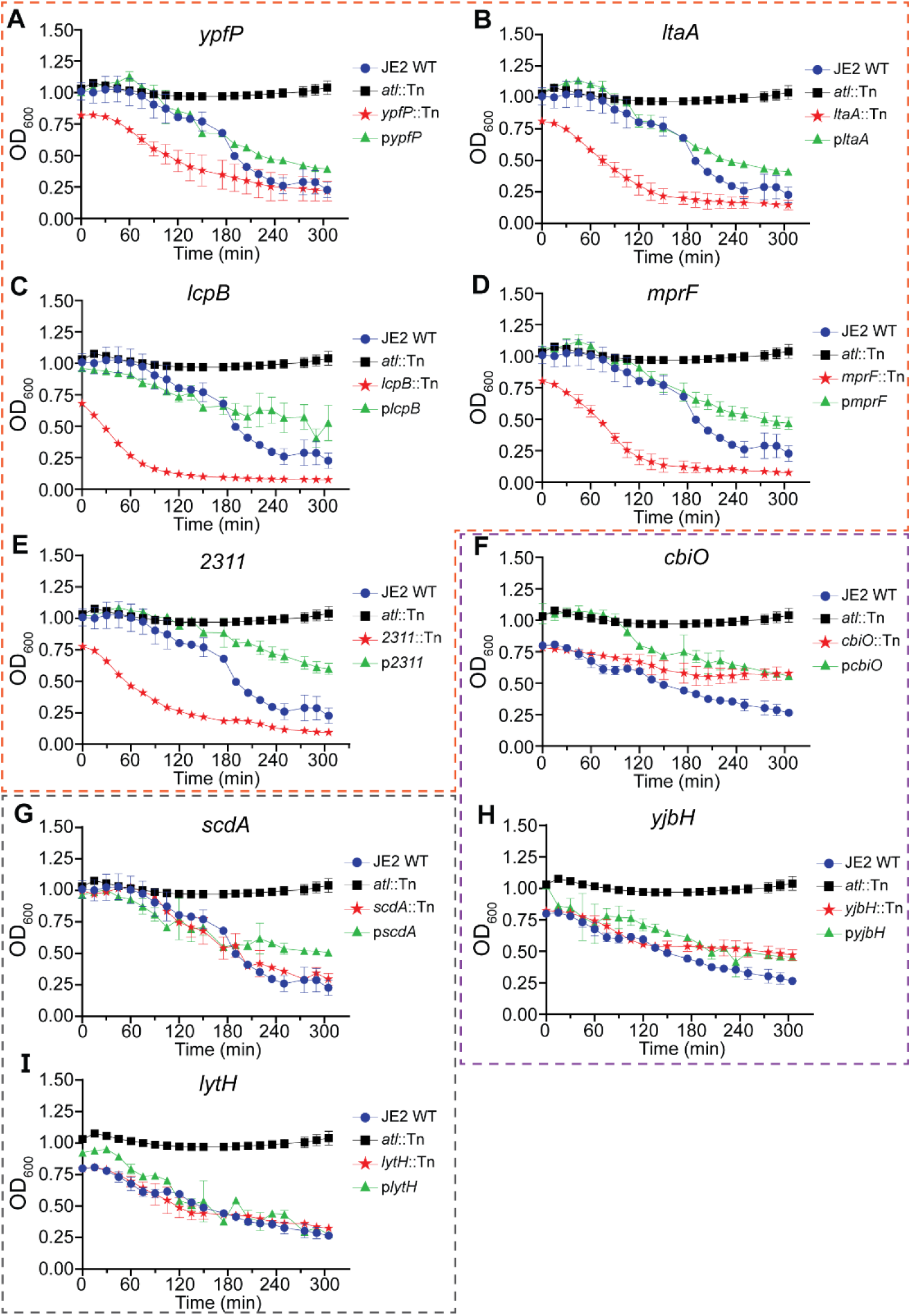
Autolysis analysis in NTML mutants. TritonX-100 mediated autolysis curves for JE2 WT (blue line), an *atlA::Tn* mutant (black line), mutant of interest (red line) and complementation strain (green line). The panels are grouped by mutants showing an increased autolysis A-E (framed by an orange dashes), a decreased autolysis F, H (framed by purple dashes) and those that showed no change in autolysis G, I (framed by gray dashes).

### Lack of WTA perturbs SpA cross-wall localization without affecting SpA septal membrane localization

The above screen from the NTML library identified several mutants that are known to be directly involved in cell envelope assembly and homeostasis, such as *ypfP, ltaA, mprF, lcpB* and *lytH*. WTA is a major component of gram-positive cell envelope. While LcpB has been implicated in ligating WTA to PG, the exact function of WTA in SpA trafficking has not been examined. To address this, we transduced a *tagO*::erm (also called *tarO*) deletion mutant deficient in WTA production to JE2, together with its complementation plasmid (Δ*tagO* comp) (59). Consistent with previous studies (60, 61), JE2Δ*tagO*::erm exhibited severe defects in cell morphology, enlarged cell size, irregular septa formation and irregular cell cluster formation (Fig. 8A). Cross-wall IF revealed that Δ*tagO*::erm disrupted SpA cross-wall localization, where SpA was mis-localized to the cell periphery of Δ*tagO*::erm (Fig. 8BC). SpA mis-localization could be complemented by expressing plasmid-borne *tagO* expression (Δ*tagO* comp). Interestingly, similar to the mutants identified from the NTML screen, Δ*tagO*::erm did not alter SpA septal membrane localization as shown by membrane-IF and quantification (Fig. 8DE). Taken together, WTA is necessary for SpA cross-wall localization. While we cannot rule out that WTA directly targets SpA, it is more likely that the *tagO* mutant also modulates the late stage SpA trafficking by altering cell wall homeostasis and cell cycle.

**Fig. 8.**
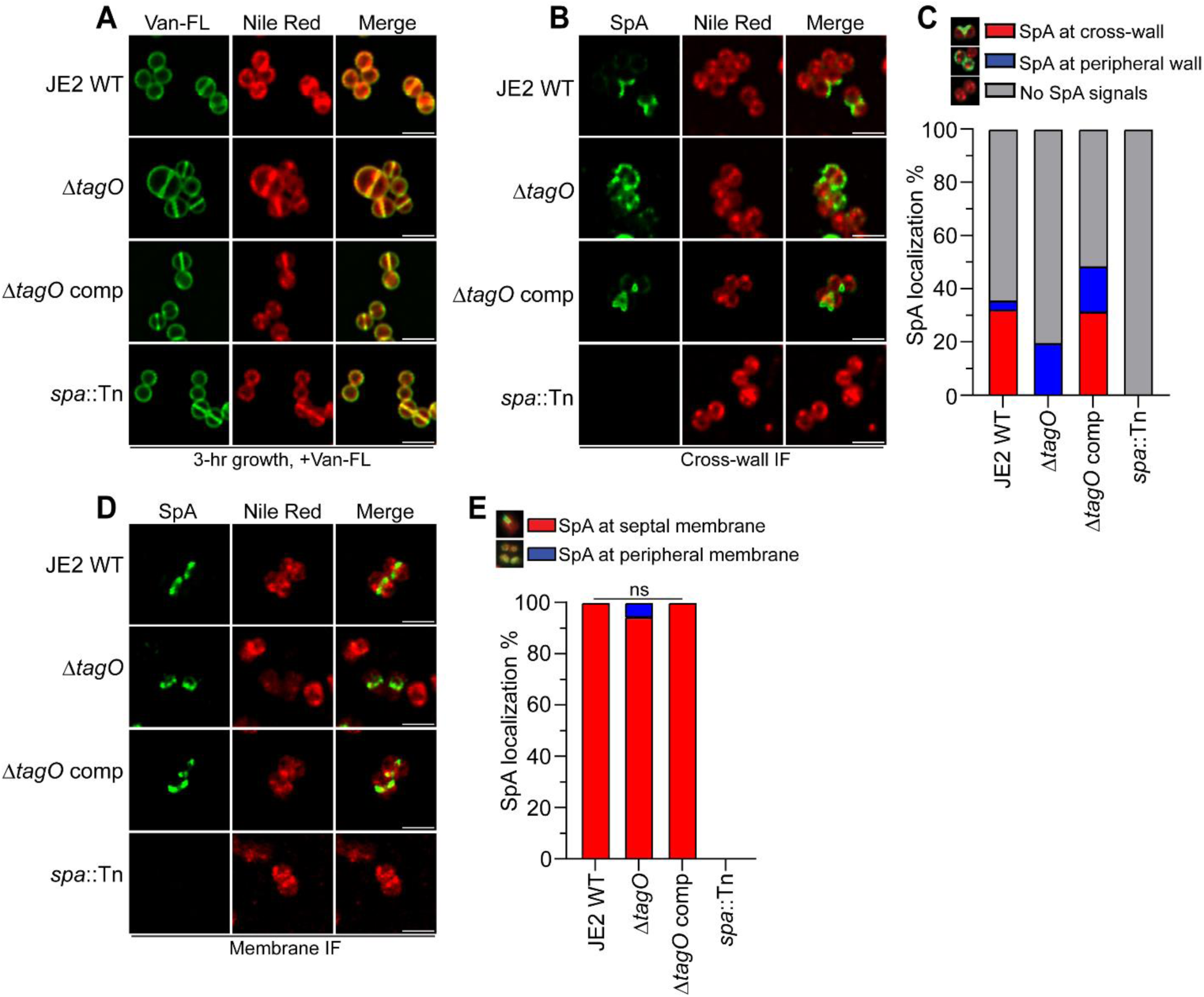
The *tagO* mutant deficient in WTA production displays SpA mis-localization and aberrant cell morphology. A) Fluorescence images showing the cell wall (Van-FL) and membrane (Nile red). B) Cross-wall IF images of JE2 WT, JE2Δ*tagO* and a *tagO* complement strain (Δ*tagO* comp). C) Quantification of SpA localization at the cross-wall (red bar), peripheral wall (blue bar) or no detectable SpA signals (gray bar) from the images represented in panel B. Mean values and standard deviation are shown in Table S6. D) Membrane IF images showing SpA membrane localization in JE2Δ*tagO*. E) Quantification of SpA localization at the septal membrane (red bar) and peripheral membrane (blue bar) from the images represented in panel D. ns = no significant difference.

## Discussion

Here we report the results of a comprehensive screen for mutants deficient in SpA cross-wall trafficking. We identified and characterized the top nine hits alongside the *tagO* mutant in terms of *spa* expression, SpA septal and cross-wall localization, cell cycle progression, cell wall synthesis and autolysis. All the mutants severely disrupted SpA cross-wall localization but did not affect SpA septal membrane localization. All the mutants displayed defects in cell morphology, cell cycle progression and dysregulated spatial distribution of PG synthesis. The shared phenotypes suggest that dysregulated PG homeostasis and impaired cell cycle are likely the mechanisms by which SpA trafficking is dysregulated. Among these ten mutants, six of them (*ypfP, ltaA, mprF, lcpB, tagO, lytH*) are known to be directly involved in the biogenesis of cell envelope structures, whereas the other four mutants (*scdA, yjbH, cbiO, 02311*) are not. Below we will discuss the possible roles of these genes in SpA trafficking and cell envelope homeostasis.

From our previous work, we showed that LtaS-mediated LTA synthesis restricts SpA septal and cross-wall localization (6, 30). LtaS localizes at the septum whereas mature LTA is more abundant at the cell periphery (10, 30, 62). The current screen identified two additional LTA pathway genes, namely *ypfP* and *ltaA*. The observation that mutants of *ypfP* and *ltaA* diminished SpA cross-wall localization is consistent with a recent study (63). However, surprisingly, mutants of *ypfP* and *ltaA* showed a notable difference compared to the *ltaS* mutant: the *ltaS* mutant dysregulates SpA cross-wall and septal membrane localization whereas the mutants of *ypfP* and *ltaA* only disrupted SpA cross-wall localization. The results suggest that YpfP and LtaA function differently compared to LtaS in term of regulating SpA trafficking. Indeed, there are several differences between LtaS and YpfP-LtaA, which may explain their functional difference. Firstly, *ltaS* is an essential gene for viability, whereas *ypfP* and *ltaA* are not.

Secondly, compared to the septal localization of LtaS, YpfP and LtaA have been shown to localize circumferentially in *S. aureus* (62). Thirdly, while *ltaS* deletion completely abolishes LTA synthesis, mutants of *ypfP* or *ltaA* produce longer LTA attached to diacylglycerol (DAG) instead of Glc2-DAG (32, 64, 65). Recent studies show that the glycolipid anchor produced by YfpP and LtaA determines LTA chain length, and the LTA chain length regulates cell size and envelope integrity (66, 67). A similar mechanism may apply to SpA spatial regulation: *ypfP* or *ltaA* mutant regulates SpA cross-wall localization due to aberrantly long LTA chain length.

MprF is a well characterized protein that synthesizes and flips lysyl-phosphatidylglycerol to the outer leaflet of cytoplasmic membrane (68, 69). The lysinylation adds positive charges to phospholipids, which leads to repulsion of cationic antimicrobial peptides (CAMPs) and resistance to CAMP-like antibiotic daptomycin (70–72). From our screen, we initially anticipated that membrane modification would impact SpA membrane localization. However, this was not the case; instead, *mprF*::Tn only disrupted SpA cross-wall localization. The unexpected results suggest that SpA mis-localization in the *mprF* mutant is not directly mediated by lysinylation of phospholipids. Interestingly, earlier work identified MprF as FmtC (factor for methicillin resistance), as it renders resistance to oxacillin (73). Cumulative literature indicates that mutations in the *mprF* gene are associated with the so called “seesaw” effect: increased daptomycin resistance sensitizes MRSA to β-lactams (74–79). The molecular basis of this effect is not fully understood. Daptomycin resistant strains often show alterations in membrane fatty acids profiles, increased membrane fluidity, and increased cell wall thickness (77, 79–81). In our previous work, we found that *dltABCD*-mediated D-alanylation of teichoic acid regulates SpA cross-wall trafficking (30). Intriguingly, dysregulation of *dlt* and *mprF* shows several similar phenotypes: both add positive charges to the cell surface and both mutants are resistant to CAMPs and daptomycin (82–84). Our work adds more shared phenotypes: both mutants diminish SpA cross-wall localization without affecting SpA septal anchoring; both have severe morphological and cell cycle defects and altered PG synthesis and autolysis. Interestingly, a recent study provides insights into possible connections between MprF and LTA: LTA modulates the cellular level and activity of cell wall-degrading enzyme LytE and MprF modulates LTA length in *B. subtilis* and *S. aureus* (85). The interactions between MprF, Dlt, LTA and SpA localization remain to be addressed in future studies.

LcpB belongs to the LytR-CpsA-Psr family proteins that function as glycopolymer ligase, which covalently attach various glycopolymers to PG (86). *S. aureus* encodes three LCP proteins, namely LcpA (MsrR), LcpB and LcpC (87). Genetic studies using single, double and triple mutants show that mutants of *lcpA* and *lcpB*, but not *lcpC* reduced WTA deposition on the cell envelope; complementation with each *lcp* gene partially restores WTA level in the triple mutant suggesting their functional redundancy (50). In vitro reconstitution experiments demonstrated that all three LCP proteins can attach WTA to PG; however synthetic lethality experiments suggest that LcpA is the major enzyme for WTA attachment to PG, LcpB contributes to WTA-related functions and LcpC is not related with WTA (88). Genetic and biochemical studies confirm that LcpC is the primary enzyme that attaches capsular polysaccharide to PG (89, 90). From our screen, *lcpC::Tn* did not affect SpA localization (data not shown). A mutant of *lcpA* is not found in the NTML. We speculated that the *lcpA* mutant or the *tagO* mutant would show similar phenotypes to *lcpB::Tn*. Indeed, a *tagO* deletion mutant exhibited SpA mis-localization as expected. The phenotype that WTA affects SpA cross-wall localization, but not septal membrane localization, can be explained as WTA is attached to PG layer; the mutant of *tagO* is severely impaired in cell division and cell separation; cell wall hydrolases are spatially dysregulated in the *tagO* mutant which may explain the imbalanced cell wall homeostasis and dysregulated SpA distribution (91, 92).

PG hydrolase-mediated cell wall turnover plays a vital role in cell wall homeostasis throughout the cell cycle (93). LytH is one of the eighteen cell wall hydrolases that *S. aureus* encodes (94). While we observed mild reduction of SpA cross-wall localization in a few other cell wall hydrolase mutants (data not shown), *lytH*::Tn showed the strongest phenotype. Interestingly, unlike other cell wall hydrolases that typically cleave cross-linked PG to separate daughter cells,

LytH has been shown to regulate cell division and growth by preferentially removing stem peptides from membrane-proximal uncross-linked PG (49). The amidase activity of LytH was shown to spatially regulate PG synthesis, whereby the *lytH* mutant decreased septal PG transpeptidation (49). LytH was also shown to form a complex with ActH protein, which is required for its amidase activity shown by in vitro reconstitution (49). The *actH* mutant displays severe cell division defects similar to the *lytH* mutant (49). In agreement with the previous study, the cell cycle and FDAA incorporation are dysregulated in *lytH::Tn* in our experiments. The *actH*::Tn exists in NTML; however, we did not observe strong SpA mis-localization in this mutant (data not shown). We also did not observe severe cell division defects nor cell size alteration in JE2 *lytH::Tn* or *actH*::Tn, which could be due to different strain backgrounds (previous work used HG003 strain). While the role of LytH and ActH in SpA trafficking needs to be examined in more detail, we propose that LytH may directly impact SpA cross-wall localization by modulating the availability of SpA anchoring sites, i.e., uncross-linked glycine bridges attached to the stem peptides, or by modulating PG transpeptidation activity required to incorporate lipid II-linked SpA into mature cell wall.

Compared to the above discussed genes, how mutants of *scdA, yjbH, cbiO* and 2*311* regulate SpA trafficking is less clear. These genes are not known to be directly involved in cell envelope biogenesis. However, all these mutants show defects in cell cycle and cell wall synthesis, implying their potential functions in cell envelope homeostasis. The *scdA* gene is located immediately downstream of WTA synthesis genes and upstream of *lytSR* two-component regulatory system that regulates autolysis (51). Earlier work demonstrates that the *scdA* mutant has severe cell division defects, increased PG crosslinking and decreased autolysis (51). ScdA has previously been implicated as a ‘repair of iron centers (RIC)’ family protein that protect iron-sulfur enzymes from nitrosative and oxidative stress (95). A recent structural study reveals that ScdA functions as a nitrite reductase that catalyzes the reduction of nitrite to nitric oxide (52).

How ScdA contributes to cell wall homeostasis is unknown. YjbH is an adapter protein that binds to the stress-response regulator Spx and enhances its proteolytic degradation by ClpXP (53, 96, 97). The *yjbH* mutant has been shown to regulate the expression of virulence genes and pathogenesis via *spx*-mediated regulation (98–101). Interestingly, mutations in *yjbH* genes are frequently associated with resistance to β-lactam antibiotics and the mechanisms are not entirely clear (102–107). Both *spx*-dependent and independent mechanisms have been reported (102, 105, 108). Moreover, the functions of YjbH seem to be pleiotropic: a recent study reports that *B. subtilis* YjbH binds to invading phage DNA via its C-terminal helix-turn-helix domain and restricts phage production to a specific subcellular site (109). In our study, we observed that the expression of *spa* is severely downregulated in *yjbH::Tn*, which is consistent with an earlier report (110). It is worth noting that three other mutants, namely *lcpB, cbiO* and *2311* also severely reduced *spa* expression. We speculate that altered *spa* expression in these mutants is due in part to cell wall stress-response. How exactly the *yjbH* mutant regulates SpA localization, cell wall homeostasis and β-lactam antibiotics resistance remains to be further studied. Little is known about the functions of the last two hits, *cbiO* and *2311.* Both mutants have been implicated in biofilm formation (111, 112). CbiO is annotated as ‘energy-coupling factor transporter ATPase’ and *‘*cobalt transporter ATP-binding subunit’ in AureoWiki (113). Ecf transporters are a subclass of ATP-binding cassette (ABC)-transporters, which mediate the uptake of essential micronutrients (114). Gene *2311* and its upstream gene *2310* are homologs of a bacterial LytTR regulatory system that regulates competence, bacteriocin production and cell death (54). The function of this system is largely unknown in *S. aureus*.

In summary, this work identified new genes required for SpA cross-wall trafficking, proper cell cycle progression and cell wall homeostasis. Our results suggest previously unknown interactions between surface protein trafficking, dynamic cell wall synthesis and turnover and teichoic acid synthesis throughout the cell cycle. Several ‘accessory’ genes that are involved in regulating cell envelope metabolism and homeostasis have been identified, which will be the subjects for future research.

## Materials and Methods

### Strains and growth conditions

The strain background for the NTML is USA300 JE2 and all transductants are in the RN4220Δ*spa* background. All *S. aureus* strains were grown on tryptic soy agar (TSA) plates and in tryptic soy broth (TSB) containing antibiotics when necessary. Broth cultures were grown at 37°C and shaken constantly at 220 rpm. The *E. coli* strains DH5α and DH5αλ*pir* were used for cloning. *E. coli* cells were grown on Miller LB agar or in Miller LB broth with shaking. Antibiotics were used at concentrations of; erythromycin 10 mg/ml (Em10), chloramphenicol 7 mg/ml (Cm7), ampicillin 100 mg/ml (Amp100) and trimethoprim 10 mg/ml (Tri10). All strains used in this study are listed in **Table S1** and all primers are listed in **Table S2.**

### Phage transduction of NTML transposon mutants

The donor strain was grown overnight in TSB. The following day cultures were diluted in heart-brain infused rich media (HiB) with 5 mM calcium chloride (CaCl_2_) and grown for 2 hours. 150 μL of *phi*85 phage lysate was added and the samples were incubated at room temperature with slight shaking overnight. The recipient strain was grown overnight in HiB with CaCl_2_. Samples were incubated for 15 minutes with 150 μL of *phi*85 phage lysate at 37°C with shaking. The samples were centrifuged for 30 seconds and resuspended in 1 mL of ice-cold 40 mM sodium citrate. Samples were pelleted and resuspended in 100 μL of sodium citrate and plated on TSA plates containing the antibiotic selection (Em10) and sodium citrate. Plates were incubated for 2 days at 37°C. Mutant-specific PCR was used to confirm transposon insertion.

To generate the Δ*tagO* mutant, we transduced an *ermB* marked deletion of *tagO* from SA113Δ*tagO*::*ermB* (strain courtesy of F. Götz lab) and selected for positive transductants on Em10 TSA plates. To generate the *tagO* complementation strain (*tagO* comp), we transduced a construct that expresses *tagO* under a constitutive promoter (pRB473-*tagO*) (construct courtesy of F. Götz lab) into the Δ*tagO* mutant generated for this study. These transductants were selected using Em10 and Cm7 TSA plates.

### Complement plasmid construction

The complementation plasmids were generated as follows: primers 626/627 were utilized to amplify the *gdpP* operon promoter (290 bp upstream of *SAUSA300_0013*). The PCR product of *gdpP* promoter and pKK30 empty vector was digested using NheI/SacI. Digested products were ligated to generate pKK30-P*_gdpP_*. Each respective gene was amplified using primers that begin at the SD sequence and amplify the gene to its stop codon. Each PCR product and pKK30-P*_gdpP_* were digested using SacI/BamHI. Digested products were ligated to generate pKK30-P*_gdpP_*-*gene* simplified as p*gene*, e.g. p*ypfP*. The exception to this process is the p*2311* construct, which contains the native promoter instead of P*_gdpP_* and both *2311* and *2310*. PCR product of *2310-2311* and pKK30 EV were digested using SacI/BamHI. Ligation reactions were transformed to DH5αλ*pir* and confirmed by sequencing. pKK30-based complement plasmid was transformed to RN4220 first and then to the respective mutant via electroporation.

### SpA cross-wall Immunofluorescence microscopy screen

We adapted and standardized the cross-wall immunofluorescence microscopy detailed in Protocol B from Scaffidi et al. 2021 (62). In brief, bacterial cells are grown overnight in TSB. The following day cells are diluted 1:100, grown to mid-exponential phase and normalized to OD_600_=0.8. Two mL of such culture was pelleted and trypsin-treated to digest existing surface proteins. Subsequently, cells are grown for precisely 20 minutes in fresh TSB with trypsin inhibitor to allow deposition of new surface proteins. Cells are immediately fixed and added onto a multi-well microscope slide. Each well is blocked with bovine serum albumin and stained with anti-rabbit SpA_KKAA_ primary antibody overnight. The following day unbound primary antibody is washed away with PBS and cells are stained with anti-rabbit IgG conjugated to Alexa Fluor 488. SpA signals are then observed via fluorescence microscopy. Each NTML strain was analyzed alongside JE2 WT, JE2 *spa*::Tn and JE2 *spa*::Tn pSP_SasD_-SpA. Images were captured by Nikon Scanning Confocal Microscope 431 ECLIPSE Ti2-E equipped with the HC PL APO 63×oil objective and analyzed in ImageJ.

Quantification of SpA localization percentage was performed on approximately 1,000 cells per strain across three biological replicates using ImageJ. Each group represents the percentage of cells with SpA at the cross-wall (red bar), SpA at the peripheral wall (blue bar) or no detectable SpA surface signal (gray bar). Unpaired t-test with Welch’s correction was performed for statistical analysis using GraphPad Prism. Values for each strain are shown in an accompanying table with standard deviations.

Quantification of SpA fluorescence intensity was performed on approximately 10-15 cells across three biological replicates using ImageJ. The method has been described previously (30). Briefly, a line was drawn perpendicular to the cross-wall at the mid-cell and the fluorescence values were plotted across the line, where mid-cell is at point ‘0’ on the x-axis.

### SpA membrane immunofluorescence microscopy

We used the protocol described as protocol C in our previous publication (38). Briefly, 2 ml of log-phase culture (OD_600_ = 0.8) were harvested and treated with trypsin. Trypsin-treated cells were fixed, washed, and resuspended in 1 ml of GTE buffer (50 mM glucose, 20 mM Tris–HCl pH 7.5, and 10 mM EDTA). 20 µg/ml of lysostaphin (AMBI) was added to each sample, and 50 µl of cell suspension was immediately applied to a poly-L-lysine-coated microscope slide and incubated for 2 min. Non-adherent cells were gently vacuumed away and any excessive liquid was aspirated. Dried slides were immediately dipped in pre-chilled methanol at −20°C for 5 min followed by dipping in pre-chilled acetone at −20°C for 30 seconds. Slides were allowed to dry at room temperature, and the samples were re-hydrated with one drop of 1X PBS for 5 min and proceed with SpA immunofluorescence microscopy as detailed above. Images were captured by Nikon Scanning Confocal Microscope 431 ECLIPSE Ti2-E equipped with the HC PL APO 63×oil objective and analyzed in ImageJ.

Quantification of SpA septal membrane localization was performed on approximately 1,000 cells per strain across three biological replicates using ImageJ. Each group represents the percentage of cells with SpA at the septal membrane (red bar) or SpA at the peripheral membrane (blue bar) in the figures. Unpaired t-test with Welch’s correction was performed for statistical analysis using GraphPad Prism.

### SpA immunoblot analysis

For all SpA Western blots, we utilized a previously established protocol (10, 30). Protein samples were prepared from cultures grown to an OD_600_ of 1.0, with supernatants (S) and cell pellets (P) separated by centrifugation. Culture supernatant was removed from cell pellets, which were then lysed with lysostaphin, and all samples underwent TCA precipitation, acetone washing, and drying before being resuspended in Tris buffer and SDS-PAGE sample buffer.

Proteins were denatured by boiling, separated on 10% SDS-PAGE gels, and transferred onto PVDF membranes. The membranes were blocked and incubated overnight with primary antibody (SpAKKAA). The following day, membranes were washed and incubated in new blocking solution containing secondary antibody with rotation at RT for 1 hour. Membranes were dipped in ECL substrate containing 3% hydrogen peroxide and then imaged on X-ray film.

Quantification of SpA abundance was performed using ImageJ gel plot tool. A box was drawn around the SpA bands associated with cell pellet (P) and another box around the secreted supernatant (S). The loading control of sortase A (SrtA) was run adjacent to the SpA blot and a box was drawn around the SrtA band. The values of SpA was divided by the SrtA band to normalize the samples to the same cell density. All blots were repeated in biological triplicate and the values shown represent the average of three blots. Unpaired t-test with Welch’s correction was performed for statistical analysis using GraphPad Prism.

### Fluorescent vancomycin (Van-FL) staining

To label bacterial cell walls with Van-FL, cells were grown overnight in TSB. The following day cultures were diluted at 1:100 in fresh TSB grown to mid-exponential phase. Cultures were washed, normalized to an OD_600_ = 1.0, chemically fixed and applied to a microscope slide. Cells were stained in the dark with 1 µg/ml BODIPY™ FL Vancomycin (Invitrogen), Hoechst 33342 DNA dye (Invitrogen) and Nile red (Sigma). Images were captured by Nikon Scanning Confocal Microscope 431 ECLIPSE Ti2-E equipped with the HC PL APO 63×oil objective and analyzed in ImageJ.

To quantify the cell cycle: at least 100 cells per sample from three independent experiments were analyzed in ImageJ. The staphylococcal cell cycle has been previously defined (55): phase 1, cells without a septum; phase 2, cells with a partial septum; phase 3, cells with a complete septum and elongated shape. Unpaired t-test with Welch’s correction was performed for statistical analysis using GraphPad Prism.

To quantify cell diameter (μm) with ImageJ; a straight line was drawn across mid-cell, and the length values were averaged across three biological experiments. Unpaired t-test with Welch’s correction was performed for statistical analysis using GraphPad Prism. Values for each strain are shown in an accompanying table with standard deviations.

### Fluorescent D-amino acid (FDAA) cell wall labeling

The method has been described in detail previously (10). Staphylococcal cultures were grown to mid-exponential phase in TSB. Cells were incubated with 500 μM HADA (TOCRIS) (blue) in TSB for 20 minutes at 37°C with shaking. Subsequently, cells were washed with PBS and incubated with 500 μM RADA (TOCRIS) (red) for another 20 minutes at 37°C while shaking.

Lastly, cells were washed with PBS and incubated with 500 μM OGDA (TOCRIS) (green) for 20 minutes. The cells were washed, fixed and applied to a slide with poly-L-lysine for microscopy. Each FDAA is fluorescent at a different wavelength, therefore, a multi-channel fluorescent microscope was used to visualize each sample. Images were captured by Nikon Scanning Confocal Microscope 431 ECLIPSE Ti2-E equipped with the HC PL APO 63×oil objective and analyzed in ImageJ.

Quantification of percentage of FDAA misincorporation was performed using ImageJ. A misincorporation was defined as aberrant localization of FDAA signal that differs from that of the WT incorporation pattern. A similar method was utilized in Zhang et al., 2025. Unpaired t-test with Welch’s correction was performed for statistical analysis using GraphPad Prism.

### Autolysis assay

To examine rates of autolysis, cells were grown to exponential phase at 37°C with shaking at 220 rpm and normalized to an OD_600_ = 1.0. Samples were centrifuged at 18,000 g for 3 minutes and pellets were washed twice with PBS. The pellets were washed once with ice-cold ddH_2_O and resuspended in PBS containing 0.05% Triton X-100. 200 μL of each sample was aliquoted into a 96-well plate and placed at 37°C with shaking for 5 hours. OD_600_ was monitored over time to measure autolysis. Experiments were performed in biological triplicate, and the average values are shown in Fig. 7.

## Acknowledgements

This work is supported by NIH/NIGMS-R35GM146993. We thank Mac Shebes for his help with the initial screen. We thank members of the Yu lab for critically reading the manuscript and providing suggestions.

---

**Fig. S1.**
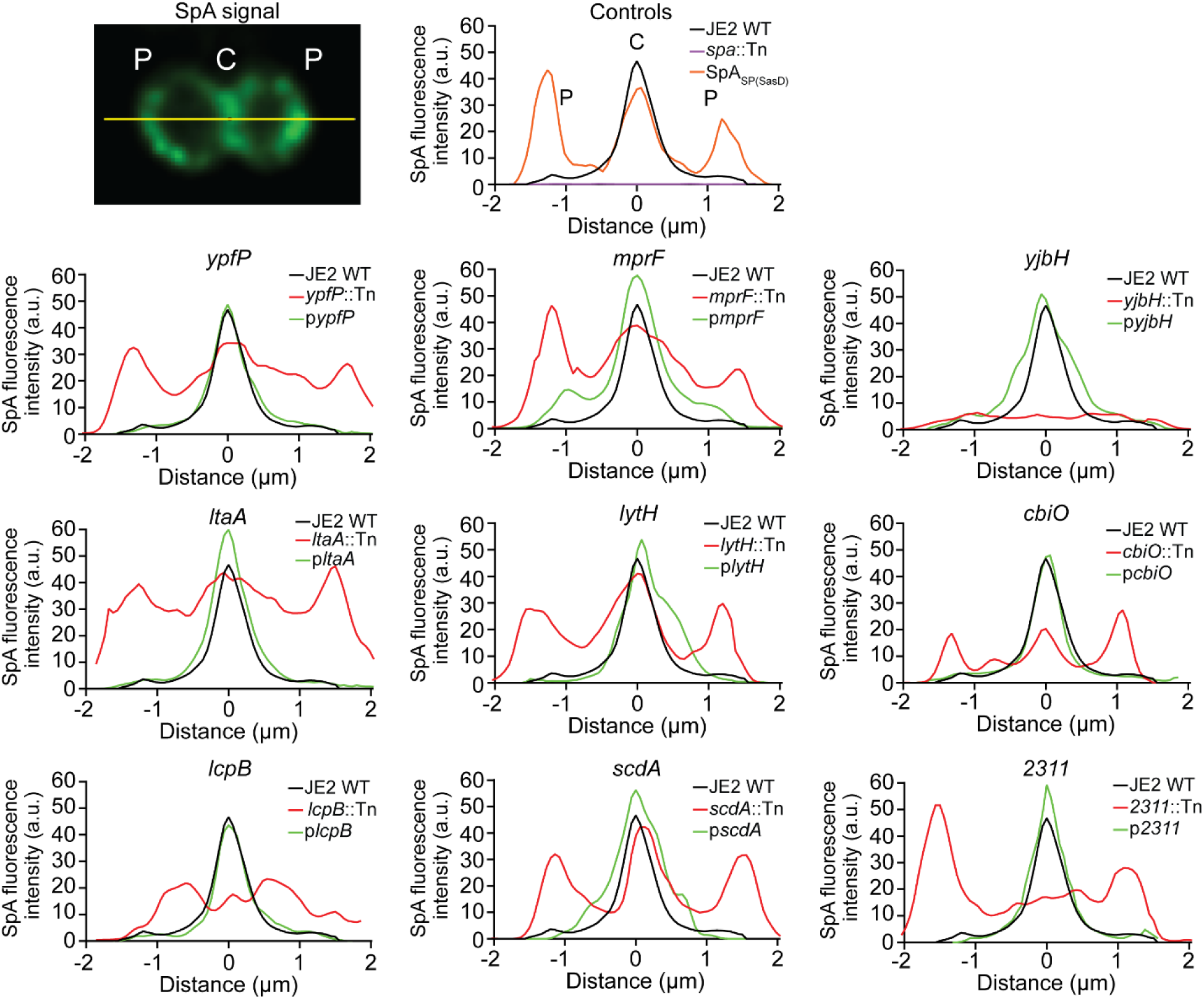
Quantificaton of SpA fluorescence intensity at cross-wall versus peripheral wall. Histogram plotting SpA fluorescence signal along a line perpendicular to the cross-wall, example image shown in upper left. The peak at distance ‘0’ represents signals at the cross-wall site (C), and the two shoulders at distance ‘-1’ and ‘1’ represent signals at peripheral wall (P). Arbitrary units (a.u.) for fluorescence intensity are used on y-axis.

**Fig. S2.**
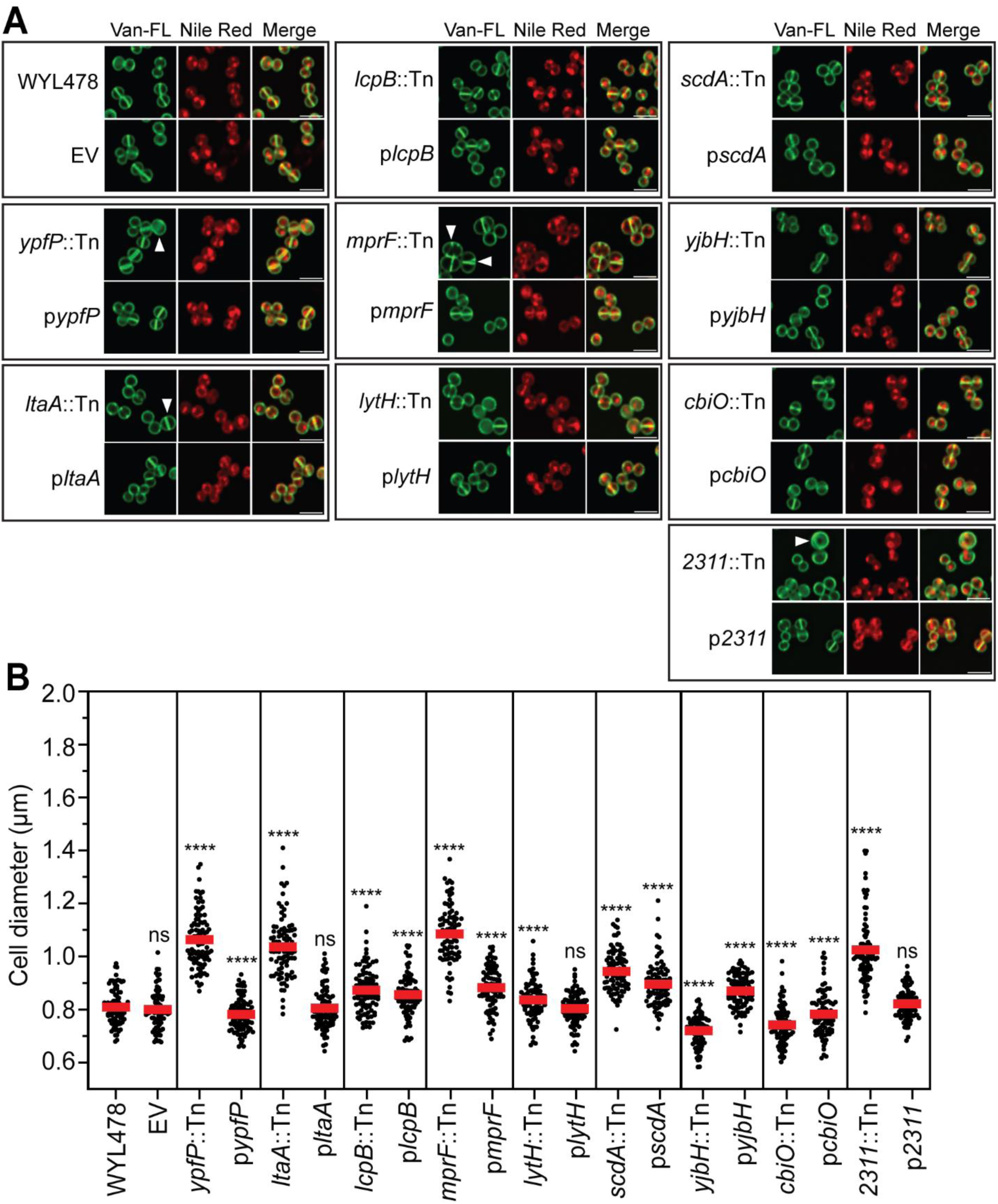
Fluorescent vancomycin (Van-FL) labeling of WYL478 transductant mutants. A) Fluorescence images showing the cell wall (Van-FL) and membrane (Nile red). White arrowheads indicate cells with aberrant morphology compared to WYL478. B) Quantification of cell diameter based on the images represented in panel A, n = 90 cells. Unpaired *t*-test with Welch’s correction was performed for statistical analysis: \**P* < 0.05; \*\**P* < 0.005; \*\*\**P* < 0.0005; \*\*\*\**P* < 0.0001.

**Fig. S3.**
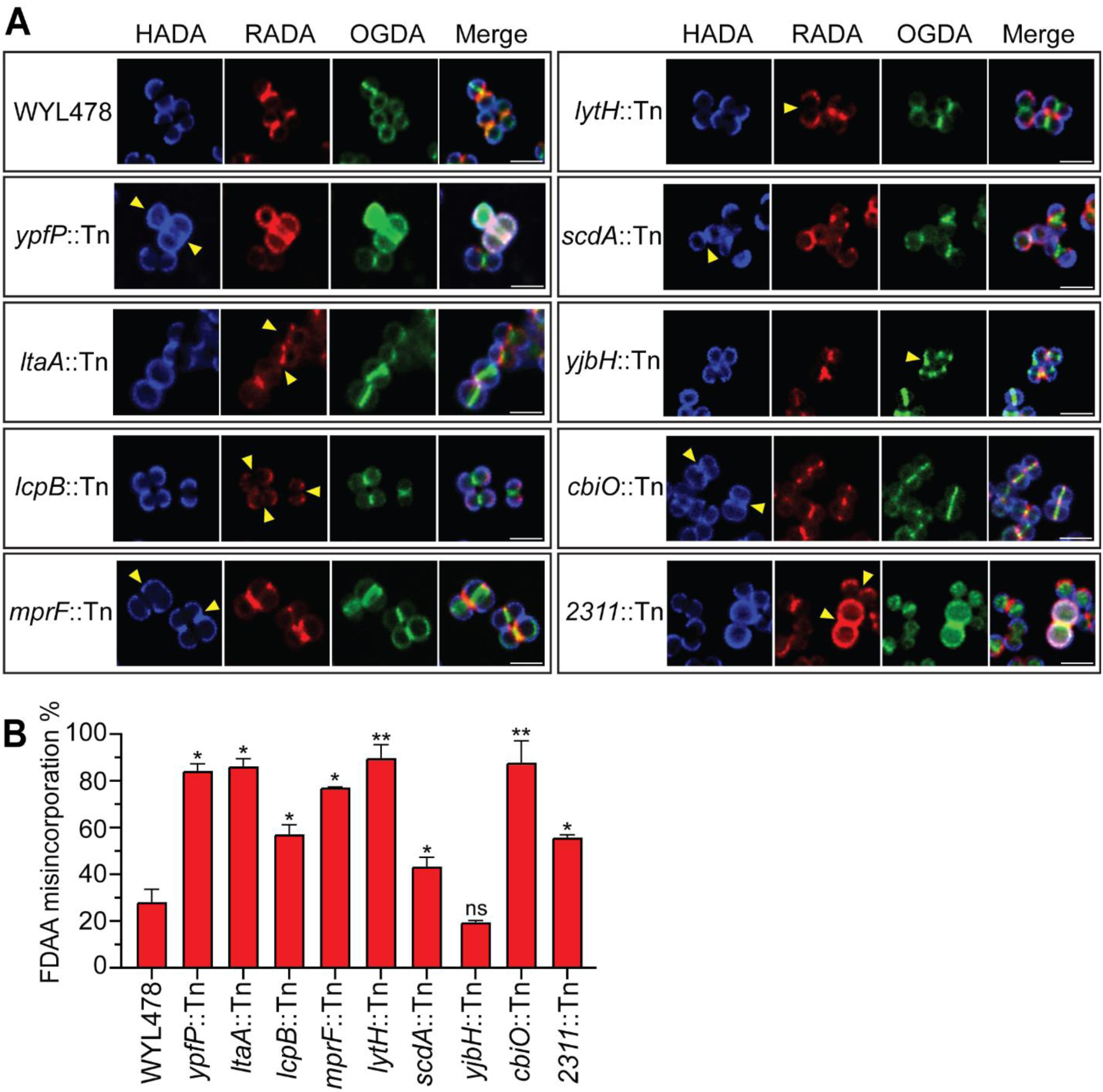
Sequential FDAAs labeling of WYL478 transductant mutants. A) Fluorescence images of cells labeled with FDAAs: HADA (blue, first), RADA (red, second) and OGDA (green, third), yellow arrows indicate aberrant FDAA localization. B) Quantification of FDAA misincorporation. Unpaired t-test with Welch’s correction was performed for statistical analysis: \**P* < 0.05, \*\**P* < 0.005. Representative images and quantification are from three independent experiments.

**Table S1.** Strains and plasmids used in this study.

| Strain or plasmid | Description | Reference or source |
| --- | --- | --- |
| *E. coli* |  |  |
| DH5α | Plasmid shuttle strain |  |
| DH5αλpir | Cloning strain for pKK30 constructs | (115) |
| *S. aureus* |  |  |
| USA300 JE2 | Wild type background for NTML | (40) |
| JE2 *spa*::Tn | Transposon mutant of protein A (*spa*) | (40) |
| JE2 *cbiO*::Tn | Transposon mutant of *cbiO* | (40) |
| JE2 *lcpB*::Tn | Transposon mutant of *lcpB* | (40) |
| JE2 *ltaA*::Tn | Transposon mutant of *ltaA* | (40) |
| JE2 *lytH*::Tn | Transposon mutant of *lytH* | (40) |
| JE2 *mprF*::Tn | Transposon mutant of *mprF* | (40) |
| JE2 2311::Tn | Transposon mutant of 2311 | (40) |
| JE2 *scdA*::Tn | Transposon mutant of *scdA* | (40) |
| JE2 *yjbH*::Tn | Transposon mutant of *yjbH* | (40) |
| JE2 *ypfP*::Tn | Transposon mutant of *ypfP* | (40) |
| JE2 Δ*tagO*::*ermB* | Deletion mutant of *tagO* | This study |
| JE2 Δ*tagO*::*ermB*<br>pRB473-*tagO* | Complementation strain for *tagO* mutant (*tagO* comp) | This study |
| WY478 | SEJ1 pCL55<sub>itet</sub>-*spa*, RN4220Δ*spa* complemented with ATc-inducible *spa* | (30) |
| WY480 | SEJ1 pCL55<sub>itet</sub>, RN4220Δ*spa* with empty vector (EV) | (30) |
| WY478 *cbiO*::Tn | Transposon mutant of *cbiO* | This study |
| WY478 *lcpB*::Tn | Transposon mutant of *lcpB* | This study |
| WY478 *ltaA*::Tn | Transposon mutant of *ltaA* | This study |
| WY478 *lytH*::Tn | Transposon mutant of *lytH* | This study |
| WY478 *mprF*::Tn | Transposon mutant of *mprF* | This study |
| WY478 2311::Tn | Transposon mutant of 2311 | This study |
| WY478 *scdA*::Tn | Transposon mutant of *scdA* | This study |
| WY478 *yjbH*::Tn | Transposon mutant of *yjbH* | This study |
| WY478 *ypfP*::Tn | Transposon mutant of *ypfP* | This study |
| pKK30 EV | Vector used for complement construction | (115) |
| pKK30-P<sub>gdpP</sub>-*cbiO* | Complementation construct for *cbiO*::Tn mutants | This study |
| pKK30-P<sub>gdpP</sub>-*lcpB* | Complementation construct for *lcpB*::Tn mutants | This study |
| pKK30-P<sub>gdpP</sub>-*ltaA* | Complementation construct for *ltaA*::Tn mutants | This study |
| pKK30-P<sub>gdpP</sub>-*lytH* | Complementation construct for *lytH*::Tn mutants | This study |
| pKK30-P<sub>gdpP</sub>-*mprF* | Complementation construct for *mprF*::Tn mutants | This study |
| pKK30-P<sub>2310/2311</sub>-*2310/2311* | Complementation construct for *2311*::Tn mutants | This study |
| pKK30-P<sub>gdpP</sub>-*scdA* | Complementation construct for *scdA*::Tn mutants | This study |
| pKK30-P<sub>gdpP</sub>-*yjbH* | Complementation construct for *yjbH*::Tn mutants | This study |
| pKK30-P<sub>gdpP</sub>-*ypfP* | Complementation construct for *ypfP*::Tn mutants | This study |

**Table S2.**
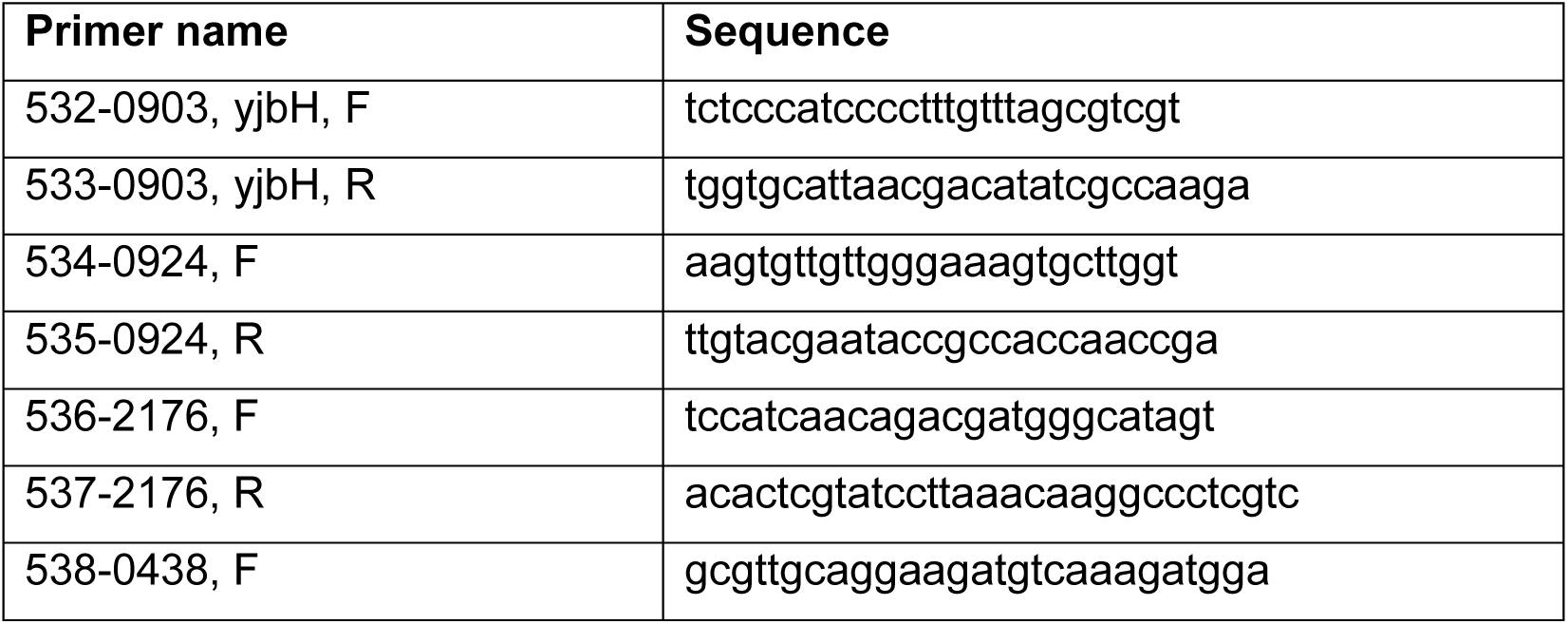
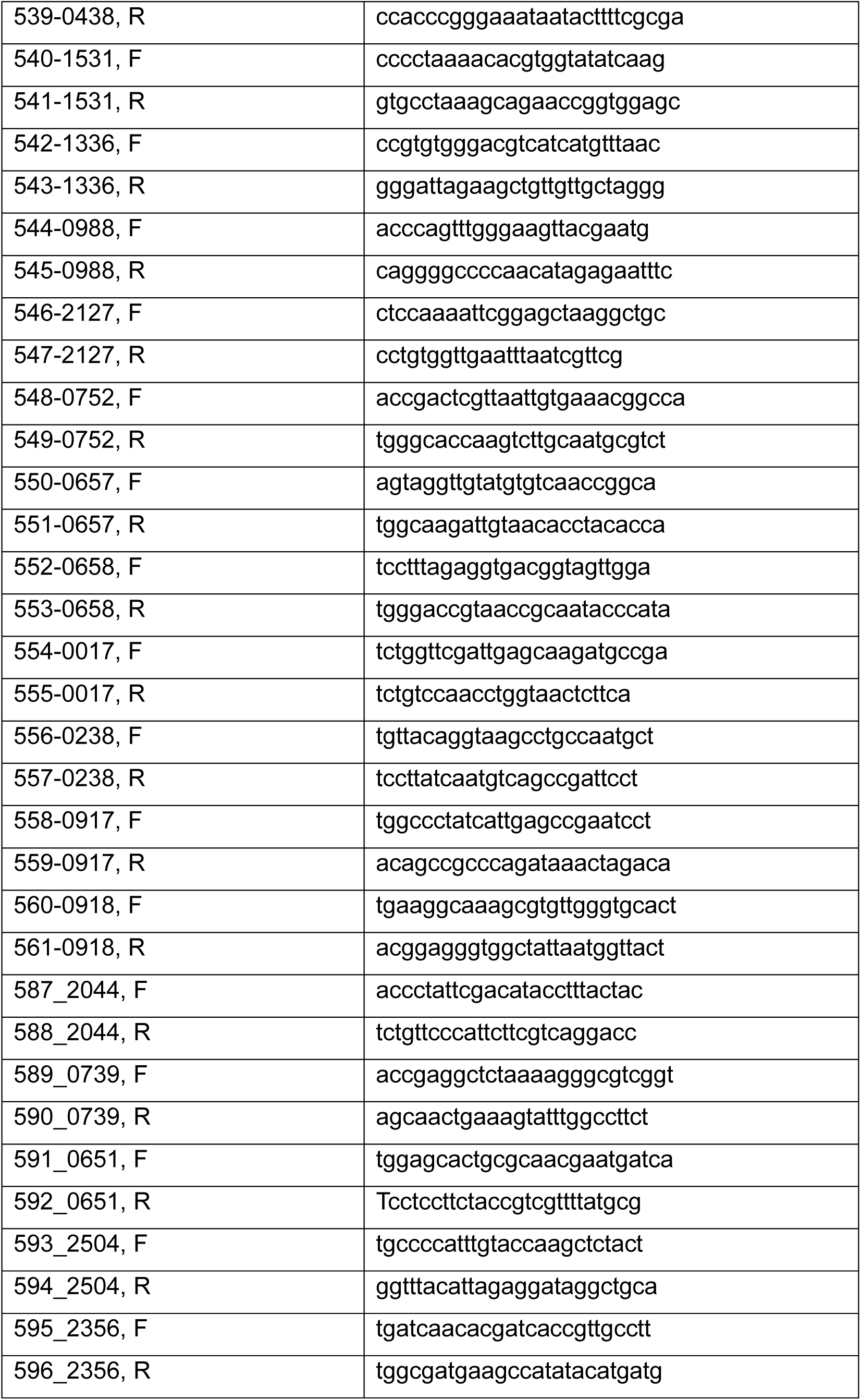
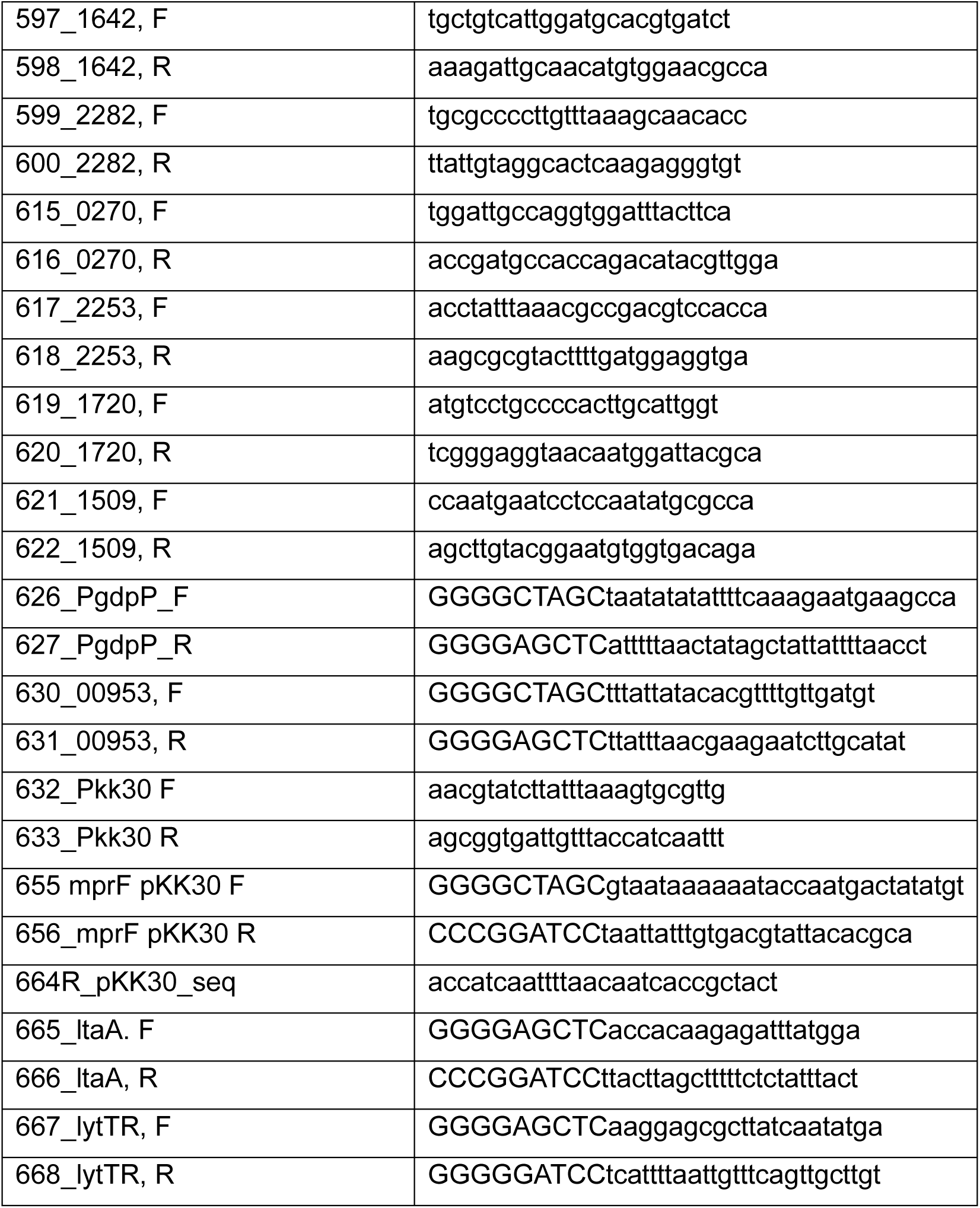
Primer list.

**Table S3.** Quantification values of SpA localization in JE2 and WYL478 background.

| Strain | Avg. % SpA at cross-wall | Avg. % SpA at peripheral wall | Avg. No SpA signal | CW p-value | PW p-value | No SpA p-value |
| --- | --- | --- | --- | --- | --- | --- |
| JE2 WT | 33.86 ± 2.7 | 3.90 ± 1.3 | 62.23 ± 3.1 |  |  |  |
| *spa::Tn* | 0.0 ± 0 | 0.0 ± 0 | 100 ± 0 | 0.0020 | 0.0358 | 0.0022 |
| SpA<sup>SP(SasD)</sup> | 1.50 ± 1.3 | 58.77 ± 8.3 | 39.73 ± 9.1 | 0.0004 | 0.0064 | 0.0394 |
| *ypfP*::Tn | 0.0 ± 0 | 5.74 ± 2.7 | 94.25 ± 2.7 | 0.0004 | <0.0001 | 0.0724 |
| *pypfP* | 32.43 ± 4.3 | 3.00 ± 2.8 | 64.55 ± 3.6 | 0.5307 | 0.2689 | 0.2856 |
| *ltaA*::Tn | 0.99 ± 1.1 | 47.93 ± 6.7 | 51.07 ± 6.1 | 0.0098 | 0.0013 | 0.0498 |
| *pltaA* | 28.48 ± 0.8 | 3.77 ± 3.3 | 67.74 ± 4.0 | 0.0302 | 0.8656 | 0.1287 |
| *lcpB*::Tn | 0.23 ± 0.4 | 10.03 ± 5.2 | 89.73 ± 5.6 | 0.0017 | 0.1737 | 0.0045 |
| *plcpB* | 26.55 ± 5.5 | 3.09 ± 3.3 | 70.34 ± 8.8 | 0.1352 | 0.7231 | 0.2462 |
| *mprF*::Tn | 6.36 ± 1.4 | 37.18 ± 5.9 | 56.45 ± 5.6 | 0.0006 | 0.0060 | 0.0676 |
| *pmprF* | 33.48 ± 2.7 | 2.56 ± 2.3 | 63.95 ± 2.3 | 0.0615 | 0.9521 | 0.1344 |
| *lytH*::Tn | 3.15 ± 3.5 | 38.80 ± 9.5 | 58.05 ± 6.0 | 0.0004 | 0.0221 | 0.3608 |
| *plytH* | 30.05 ± 5.2 | 1.47 ± 1.8 | 68.46 ± 23.6 | 0.3387 | 0.1359 | 0.0869 |
| *scdA*::Tn | 0.43 ± 0.7 | 0.36 ± 0.6 | 99.20 ± 1.4 | 0.0006 | 0.0078 | 0.2101 |
| *pscdA* | 32.32 ± 4.3 | 4.80 ± 3.2 | 62.87 ± 2.6 | 0.8734 | 0.4419 | 0.4868 |
| *yjbH*::Tn | 0.0 ± 0 | 6.91 ± 2.5 | 93.08 ± 2.5 | 0.0011 | 0.0268 | 0.0005 |
| *pyjbH* | 27.50 ± 7.8 | 1.29 ± 1.1 | 71.20 ± 7.9 | 0.6325 | 0.6848 | 0.7972 |
| *cbiO*::Tn | 4.16 ± 3.4 | 39.05 ± 1.7 | 56.78 ± 1.9 | 0.0020 | 0.1644 | 0.0002 |
| *pcbiO* | 32.55 ± 1.9 | 1.99 ± 2.1 | 65.44 ± 3.3 | 0.2894 | 0.0604 | 0.1776 |
| *2311*::Tn | 1.9 ± 0.8 | 27.81 ± 4.3 | 70.28 ± 3.5 | 0.0020 | 0.3701 | 0.0002 |
| *p2311* | 33.56 ± 3.7 | 1.51 ± 1.7 | 64.92 ± 3.0 | 0.6558 | 0.6494 | 0.4482 |
| **Strain** | **Avg. % SpA at cross-wall** | **Avg. % SpA at peripheral wall** | **Avg. No SpA signal** | **CW p-value** | **PW p-value** | **No SpA p-value** |
| WYL478 | 33.86 ± 1.4 | 6.55 ± 2.3 | 59.58 ± 1.6 |  |  |  |
| EV | 0.0 ± 0 | 0.0 ± 0 | 100 ± 0 | 0.0005 | 0.0405 | 0.0005 |
| SpA<sub>SP(SasD)</sub> | 0.92 ± 1.6 | 43.32 ± 5.9 | 55.75 ± 6.9 | <0.0001 | 0.0036 | 0.4398 |
| *ypfP*::Tn | 1.61 ± 1.4 | 44.23 ± 6.3 | 54.15 ± 6.3 | <0.0001 | 0.0044 | 0.2735 |
| *pypfP* | 27.94 ± 7.9 | 1.80 ± 1.6 | 70.24 ± 7.3 | 0.3249 | 0.0523 | 0.1210 |
| *ltaA*::Tn | 10.19 ± 6.2 | 44.23 ± 6.9 | 45.57 ± 7.3 | 0.0180 | 0.0062 | 0.0731 |
| *pltaA* | 26.35 ± 8.9 | 4.73 ± 4.6 | 69.01 ± 8.5 | 0.2817 | 0.5843 | 0.1905 |
| *lcpB*::Tn | 3.13 ± 1.6 | 36.43 ± 7.1 | 60.43 ± 6.4 | <0.0001 | 0.0117 | 0.8410 |
| *plcpB* | 25.77 ± 2.1 | 0.69 ± 1.2 | 73.52 ± 3.3 | 0.0083 | 0.0316 | 0.0082 |
| *mprF*::Tn | 0.0 ± 0 | 51.43 ± 3.6 | 48.56 ± 3.6 | 0.0005 | 0.0002 | 0.0208 |
| *pmprF* | 28.44 ± 7.4 | 3.54 ± 3.1 | 68.33 ± 8.1 | 0.3335 | 0.2596 | 0.1983 |
| *lytH*::Tn | 4.34 ± 2.3 | 51.83 ± 11.2 | 43.81 ± 9.5 | 0.0002 | 0.0165 | 0.0997 |
| *plytH* | 20.17 ± 7.0 | 0.0 ± 0 | 79.82 ± 7.0 | 0.0724 | 0.0405 | 0.0323 |
| *scdA::Tn* | 3.73 ± 4.6 | 28.38 ± 12.6 | 67.88 ± 14.8 | 0.0045 | 0.0913 | 0.4336 |
| *p<sub>scdA</sub>* | 19.07 ± 3.6 | 1.40 ± 2.4 | 79.52 ± 2.5 | 0.0116 | 0.0577 | 0.0007 |
| *yjbH::Tn* | 0.0 ± 0 | 0.0 ± 0 | 100 ± 0 | 0.0005 | 0.0405 | 0.0005 |
| *pyjbH* | 28.86 ± 11.0 | 2.82 ± 2.5 | 68.34 ± 8.5 | 0.5139 | 0.1368 | 0.2140 |
| *cbiO::Tn* | 2.19 ± 0.2 | 41.36 ± 2.2 | 56.44 ± 2.2 | 0.0005 | <0.0001 | 0.1281 |
| *p<sub>cbiO</sub>* | 31.74 ± 15.1 | 6.35 ± 5.0 | 62.01 ± 16.7 | 0.8300 | 0.9562 | 0.8250 |
| *2311::Tn* | 3.18 ± 0.7 | 36.86 ± 2.3 | 59.96 ± 2.1 | <0.0001 | <0.0001 | 0.8184 |
| *p2311* | 26.65 ± 3.4 | 3.63 ± 2.2 | 69.92 ± 2.7 | 0.0534 | 0.1894 | 0.0084 |

**Table S4.** Quantification values of cell diameter in JE2 and WYL478 background.

| Strain | Average diameter (μm) | p-value | Strain | Average diameter (μm) | p-value |
| --- | --- | --- | --- | --- | --- |
| JE2 WT | 0.848 ± 0.05 |  | WYL478 | 0.809 ± 0.03 |  |
| *spa::Tn* | 0.851 ± 0.04 | 0.8169 | EV | 0.799 ± 0.02 | 0.3846 |
| *ypfP::Tn* | 0.981 ± 0.04 | <0.0001 | *ypfP::Tn* | 1.064 ± 0.02 | <0.0001 |
| *pypfP* | 0.852 ± 0.06 | 0.5048 | *pypfP* | 0.781 ± 0.01 | 0.0044 |
| *ltaA::Tn* | 1.00 ± 0.04 | <0.0001 | *ltaA::Tn* | 1.034 ± 0.06 | <0.0001 |
| *pltaA* | 0.864 ± 0.07 | 0.1559 | *pltaA* | 0.806 ± 0.04 | 0.7095 |
| *lcpB::Tn* | 0.898 ± 0.01 | <0.0001 | *lcpB::Tn* | 0.873 ± 0.03 | <0.0001 |
| *plcpB* | 0.856 ± 0.05 | 0.4241 | *plcpB* | 0.854 ± 0.03 | <0.0001 |
| *mprF::Tn* | 1.00 ± 0.02 | <0.0001 | *mprF::Tn* | 1.085 ± 0.01 | <0.0001 |
| *pmprF* | 0.841 ± 0.04 | 0.6700 | *pmprF* | 0.881 ± 0.05 | <0.0001 |
| *lytH::Tn* | 0.841 ± 0.02 | 0.6969 | *lytH::Tn* | 0.836 ± 0.02 | 0.0080 |
| *plytH* | 0.849 ± 0.02 | 0.8165 | *plytH* | 0.803 ± 0.01 | 0.5529 |
| *scdA::Tn* | 0.902 ± 0.05 | <0.0001 | *scdA::Tn* | 0.943 ± 0.02 | <0.0001 |
| *p<sub>scdA</sub>* | 0.872 ± 0.01 | 0.0212 | *p<sub>scdA</sub>* | 0.894 ± 0.05 | <0.0001 |
| *yjbH::Tn* | 0.742 ± 0.02 | <0.0001 | *yjbH::Tn* | 0.720 ± 0.02 | <0.0001 |
| *pyjbH* | 0.880 ± 0.03 | 0.0010 | *pyjbH* | 0.869 ± 0.01 | <0.0001 |
| *cbiO::Tn* | 0.751 ± 0.01 | <0.0001 | *cbiO::Tn* | 0.741 ± 0.01 | <0.0001 |
| *p<sub>cbiO</sub>* | 0.851 ± 0.07 | 0.7749 | *p<sub>cbiO</sub>* | 0.781 ± 0.03 | 0.0171 |
| *2311::Tn* | 1.09 ± 0.03 | <0.0001 | *2311::Tn* | 1.025 ± 0.01 | <0.0001 |
| *p2311* | 0.859 ± 0.01 | 0.2235 | *p2311* | 0.822 ± 0.02 | 0.1743 |

**Table S5.** Quantification values of cell cycle analysis in JE2 background.

| Strain | phase 1 | phase 2 | phase 3 | phase 1<br>p-value | phase 2<br>p-value | phase 3<br>p-value |
| --- | --- | --- | --- | --- | --- | --- |
| JE2 WT | 67.63 ± 1.34 | 13.12 ± 0.61 | 19.25 ± 0.92 |  |  |  |
| *ypfP*::Tn | 68.01 ± 1.07 | 21.98 ± 2.85 | 20.03 ± 1.52 | 0.0019 | 0.0282 | 0.4995 |
| *pypfP* | 57.99 ± 1.71 | 13.59 ± 1.41 | 18.39 ± 2.01 | 0.7178 | 0.6329 | 0.5520 |
| *ltaA*::Tn | 73.42 ± 4.41 | 29.71 ± 3.10 | 12.95 ± 1.06 | 0.0307 | 0.0205 | 0.0010 |
| *pltaA* | 57.34 ± 3.65 | 10.88 ± 1.75 | 15.70 ± 2.95 | 0.1415 | 0.1465 | 0.1637 |
| *lcpB*::Tn | 72.22 ± 4.27 | 18.32 ± 2.20 | 12.66 ± 3.03 | 0.6568 | 0.0464 | 0.0535 |
| *plcpB* | 69.02 ± 4.59 | 12.12 ± 1.89 | 15.66 ± 2.75 | 0.1967 | 0.4625 | 0.1417 |
| *mprF*::Tn | 65.68 ± 1.99 | 23.36 ± 2.78 | 14.39 ± 2.22 | 0.0071 | 0.0197 | 0.0474 |
| *pmprF* | 62.25 ± 0.83 | 11.69 ± 0.96 | 22.36 ± 1.26 | 0.2420 | 0.1660 | 0.0298 |
| *lytH*::Tn | 66.87 ± 2.25 | 18.82 ± 3.35 | 16.24 ± 2.21 | 0.0565 | 0.0045 | 0.1017 |
| *plytH* | 64.93 ± 3.10 | 13.79 ± 2.73 | 19.33 ± 2.40 | 0.6489 | 0.7128 | 0.9617 |
| *scdA*::Tn | 69.20 ± 3.24 | 28.78 ± 2.01 | 17.16 ± 4.12 | 0.0451 | 0.0030 | 0.4739 |
| *pscdA* | 54.06 ± 5.56 | 12.65 ± 2.47 | 18.15 ± 1.85 | 0.5001 | 0.7758 | 0.4253 |
| *yjbH*::Tn | 67.53 ± 3.78 | 25.21 ± 5.53 | 12.73 ± 2.59 | 0.0042 | 0.0002 | 0.0431 |
| *pyjbH* | 62.06 ± 3.78 | 13.37 ± 3.03 | 19.10 ± 1.11 | 0.9693 | 0.8995 | 0.8625 |
| *cbiO*::Tn | 67.46 ± 1.65 | 20.28 ± 4.29 | 16.95 ± 2.54 | 0.1970 | 0.0249 | 0.2622 |
| *pcbiO* | 62.77 ± 6.80 | 12.73 ± 2.60 | 19.81 ± 1.70 | 0.8985 | 0.8235 | 0.6533 |
| *2311*::Tn | 55.00 ± 7.50 | 27.77 ± 7.09 | 17.23 ± 3.62 | 0.0011 | 0.0082 | 0.1005 |
| *p2311* | 69.85 ± 4.80 | 11.81 ± 1.68 | 18.34 ± 3.20 | 0.5114 | 0.3102 | 0.6760 |

**Table S6.**
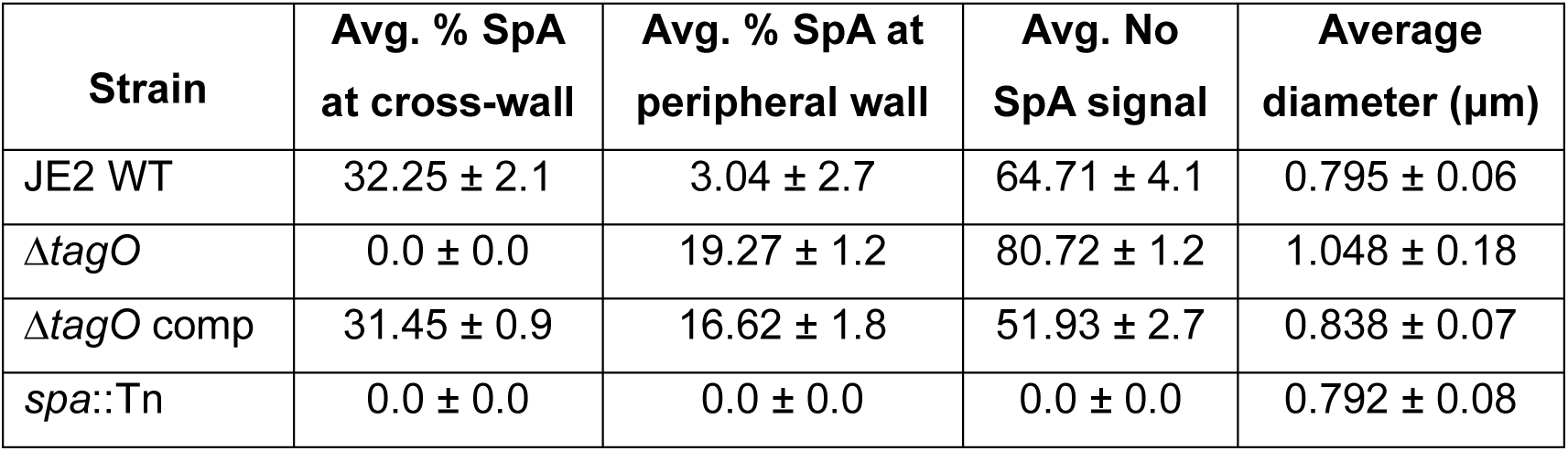
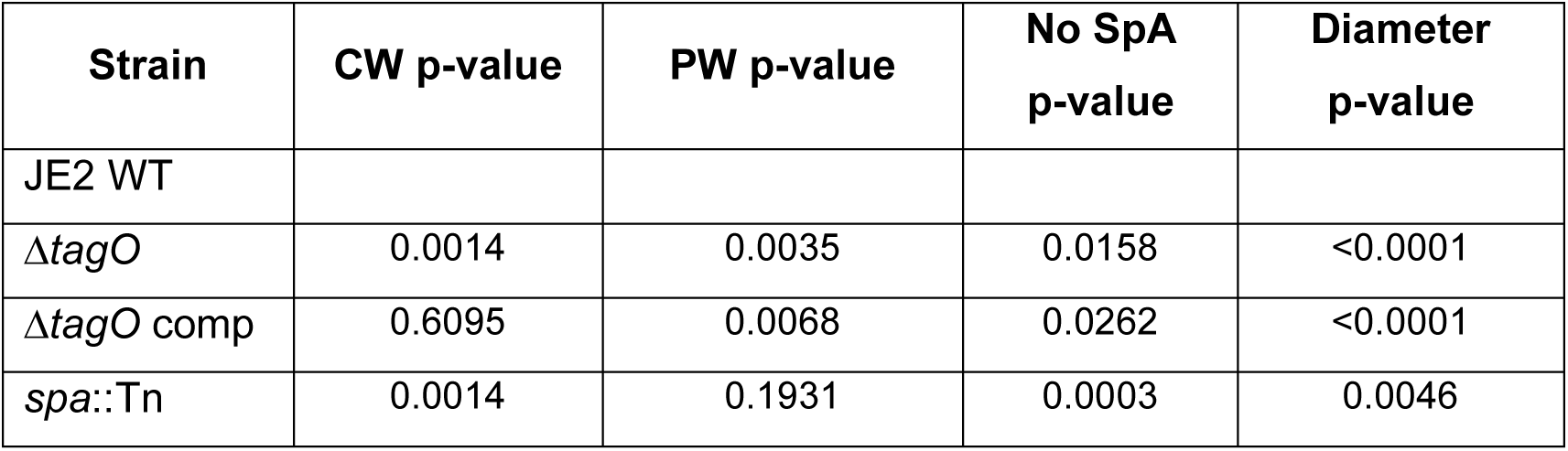
Quantification values of SpA localization and cell diameter of the *ΔtagO* mutant.

| Strain | Avg. % SpA<br>at cross-wall | Avg. % SpA at<br>peripheral wall | Avg. No<br>SpA signal | Average<br>diameter (μm) |
| --- | --- | --- | --- | --- |
| JE2 WT | 32.25 ± 2.1 | 3.04 ± 2.7 | 64.71 ± 4.1 | 0.795 ± 0.06 |
| $\Delta tagO$ | 0.0 ± 0.0 | 19.27 ± 1.2 | 80.72 ± 1.2 | 1.048 ± 0.18 |
| $\Delta tagO$ comp | 31.45 ± 0.9 | 16.62 ± 1.8 | 51.93 ± 2.7 | 0.838 ± 0.07 |
| *spa*::Tn | 0.0 ± 0.0 | 0.0 ± 0.0 | 0.0 ± 0.0 | 0.792 ± 0.08 |

| Strain | CW p-value | PW p-value | No SpA p-value | Diameter p-value |
| --- | --- | --- | --- | --- |
| JE2 WT |  |  |  |  |
| $\Delta tagO$ | 0.0014 | 0.0035 | 0.0158 | <0.0001 |
| $\Delta tagO$ comp | 0.6095 | 0.0068 | 0.0262 | <0.0001 |
| *spa::Tn* | 0.0014 | 0.1931 | 0.0003 | 0.0046 |

